# A bird’s-eye view: exploration of the flavin-containing monooxygenase (FMO) superfamily in common wheat

**DOI:** 10.1101/2024.01.16.575770

**Authors:** Sherry Sun, Guus Bakkeren

**Affiliations:** The University of British Columbia, Department of Botany, 6270 University Blvd, Vancouver, BC V6T 1Z4, Canada; Agriculture and Agri-Food Canada, Summerland Research & Development Centre, Summerland, BC V0H 1Z0, Canada

**Author notes:** Corresponding author: Guus Bakkeren, *Email:.

**Keywords:** disease resistance, FMO, gene family, phylogenetics, wheat

## Abstract

The Flavin Monooxygenase (*FMO*) gene superfamily in plants is involved in various processes most widely documented for its involvement in auxin biosynthesis, specialized metabolite biosynthesis, and plant microbial defense signalling. The roles of FMOs in defense signaling and disease resistance have recently come into focus as they may present opportunities to increase immune responses in plants including leading to systemic acquired resistance, but are not well characterized. We present a comprehensive catalogue of *FMO*s found in genomes across vascular plants and explore, in depth, 170 wheat *TaFMO* genes for sequence architecture, *cis-* acting regulatory elements, and changes due to Transposable Element insertions. A molecular phylogeny separates TaFMOs into three clades (A, B, and C) for which we further report gene duplication patterns, and differential rates of homoeologue expansion and retention among TaFMO subclades. We discuss Clade B TaFMOs where gene expansion is similarly seen in other cereal genomes. Transcriptome data from various studies point towards involvement of subclade B2 *TaFMO*s in disease responses against both biotrophic and necrotrophic pathogens, substantiated by promoter element analysis. We hypothesize that certain TaFMOs are responsive to both abiotic and biotic stresses, providing potential targets for enhancing disease resistance, plant yield and other important agronomic traits. Altogether, FMOs in wheat and other crop plants present an untapped resource to be exploited for improving the quality of crops.

## Introduction

Flavin-containing monooxygenases (FMOs; EC 1.14.13.8) constitute a ubiquitous class of ancient, highly-conserved enzymes that have been well-characterized in bacteria, fungi, and insects, catalyzing the transformations of many molecules across all domains of life (Eswaramoorthy et al., 2006; Huijbers et al., 2014; Mascotti et al., 2015). FMOs are dimeric in nature, with Rossman folds in a bi-lateral distribution containing FAD- and NAD(P)H-binding domains (Rossner et al., 2017; Schlaich, 2007; Thodberg & Jakobsen Neilson, 2020a). Typically, FMOs act on small, nucleophilic substrates that contain either a sulfur (S) or nitrogen (N) atom, resulting in the oxygenation of a wide range of compounds; this process often aides in the detoxification of xenobiotic compounds in many organisms (Eswaramoorthy et al., 2006). While FMOs are well characterized in bacteria, fungi, and insects, less is known of many of the FMOs in vascular plants, despite abundant putative plant FMOs having been observed (Schlaich, 2007).

Multiple studies from the last two decades indicate that a distinct subset of plant FMOs have evolved to participate in biosynthesis of auxin, a crucial hormone for plant growth (Kendrew, 2001a; Schlaich, 2007; Yamamoto et al., 2007; Y. Zhao, 2014). This specialized group of plant FMOs has been coined ‘YUCCA’, named after the phenotype of a dominant *Arabidopsis thaliana fmo* deletion mutant resembling the yucca plant (Y. Zhao et al., 2001) and many of which have been functionally characterized in this model plant (Cao et al., 2019; Qin et al., 2020). Due to the significance auxin biosynthesis has on plant growth, development, and yield in crop plants, many studies since then have focused on identifying and characterising the orthologous ‘YUCCA’ FMOs across a range of commercially valuable plant species, such as apple, peach, soybean, and rice (Luo et al., 2022; Qin et al., 2020; Song et al., 2020; Yamamoto et al., 2007; Zhang et al., 2022). Additionally, a subset of FMOs in *A. thaliana* have been characterized for their activity in the *S*-oxygenation of sidechains in various glucosinolate (GSL) compounds (Li et al., 2011). GSLs are potent, specialized defense metabolites found exclusively in the order Brassicales known to play a major role against insect herbivory (Cang et al., 2018; Kong et al., 2016). Grouped as “FMO-GSOXs”, these FMOs, which participate in processing metabolites, play a significant defensive role in plants that produce GSLs. Of the 29 total identified FMOs in *A. thaliana*, eleven are annotated to be YUCCAs, twelve are purportedly FMO-GS-OXs, one has been reported to be crucial for systemic acquired resistance (SAR) against microbial pathogens (Czarnocka et al., 2020; Hartmann et al., 2018), and five do not presently have designated functions. A recent review (Mitchell & Weng, 2019) provided a brief synopsis of these three groups of plant FMOs. Together these studies reveal that FMOs play diverse biological responses in plants, much of which have yet to be characterised.

Outside *A. thaliana*, much less is known about non-YUCCA FMOs, particularly in cereal crops such as *Triticum aestivum* (common bread wheat), an essential food source for over 50% of the world’s population (Curtis & Halford, 2014). *Yang et al.* (Yang et al., 2021) focussed on ‘YUCCA’ genes in wheat and identified 63 *TaYUCCA FMO* genes which they assigned to six subclades in a gene genealogy comparison; for a subset of these genes, they analyzed transcriptional activity using public transcriptomic data in an attempt to reveal function. Gaba et al. (2023) recently surveyed the genomes of ten different wild and cultivated rice species for *FMO* genes and revealed how little is presently known about FMOs outside the YUCCA group. Their study emphasized the need for more in-depth studies of other FMOs in crop plants.

Here, we address the extent to which *FMOs* have expanded across vascular plants by surveying comprehensively *FMO*-encoding genes across a broad range of plant taxa. We identified 170 likely wheat *FMO*s (*TaFMO*s) in the cultivar ‘Chinese Spring’ (the fully-sequenced reference genome, RefSeq v2.1), and sought to expand knowledge of the evolution, distribution, gene expansions, and potential functional diversity for different groups of *TaFMO* genes by using thorough phylogenetic, gene genealogy and transcriptome analyses using data obtained from publicly available studies. In addition, we take into account existing literature pertaining to *TaYUCCAs* (Li et al., 2014; Yang et al., 2021) and present a more cohesive picture of the biological roles that *TaFMO*s may play across a broad range of conditions, with a particular focus on various pathogen defense responses.

## Materials and Methods

### Genome-wide identification of FMO genes

We obtained a protein-family hidden Markov model (HMM) profile for the FMO-like family (PF00743 v22) from the Protein Family (Pfam) database (https://pfam.xfam.org/), and conducted a local HMM search (HMMER v3.3.2; http://hmmer.org/) with significance parameters of *E-*value < 1e^-2^ against the International Wheat Genome Sequencing Consortium (IWGSC) fully-sequenced reference genomes for version RefSeqv2.1, to identify all putative wheat *flavin-containing monooxygenase* genes (*TaFMOs*) for the cultivar ‘Chinese Spring’. Results from HMMER were further consolidated with the use of query-based searches in both the online database Wheat Proteome (Duncan et al., 2017) with the search term ‘flavin-containing monooxygenase’ and the search term ‘flavin monooxygenase’ in the UniProt database (https://www.uniprot.org/). Similarly, all *FMO* candidate genes from other plant species spanning a vast range of green plants and algae, were found through a HMM-based search. Table 1 lists the total number of FMOs per plant species. We used the summary of relationships of major angiosperm lineages available at NCBI Common Tree builder (https://www.ncbi.nlm.nih.gov/Taxonomy/CommonTree/wwwcmt.cgi) to display the distribution of FMOs in vascular plants (Figure 1; for sources of sequence and protein data and gene search details, see Table S1, S2; Appendix S1, Figure S1).

**Figure 1.**
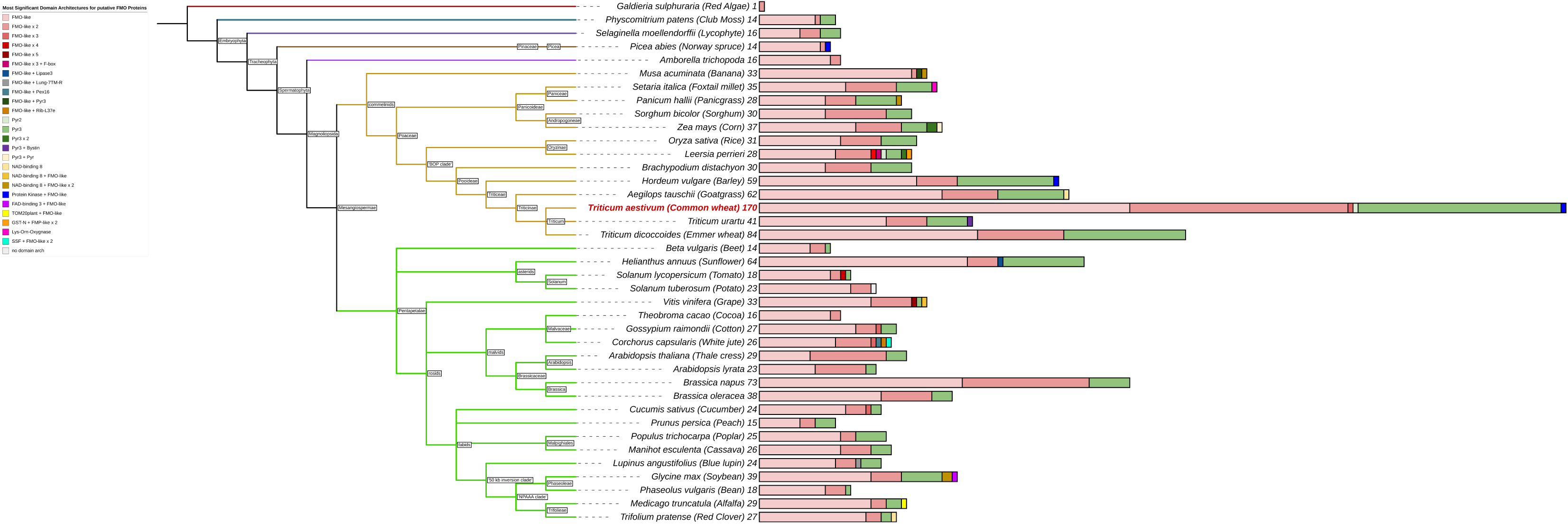
Total number of predicted FMOs and their main domain architectures as reported from Hidden Markov Modelling (HMM). Branch colours indicate major subdivisions amongst vascular plants. Total numbers of FMOs are reported as a number at the tips of each bar, and multivariate bars shows colours corresponding to the proportion of FMOs for each of the various kinds of domain architectures seen in the legend.

**Table 1.** List of plant species in FMO search (HMM profile PF00743) and number of total FMOs found.

| Species | Common Name | FMO total number |
| --- | --- | --- |
| *Aegilops tauschii* | Goatgrass | 62 |
| *Amborella trichopoda* | N/A | 16 |
| *Arabidopsis lyrata* | Lyre-leaved Rock Cress | 23 |
| *Arabidopsis thaliana* | Thale Cress | 29 |
| *Beta vulgaris* | Beet | 14 |
| *Brachypodium distachyon* | Stiff Brome | 30 |
| *Brassica napus* | Rapeseed | 73 |
| *Brassica oleracea* | Cabbage/Broccoli/etc. | 38 |
| *Corchorus capsularis* | White Jute | 26 |
| *Cucumis sativus* | Cucumber | 24 |
| *Galdieria sulphuraria* | Red Microalga | 1 |
| *Glycine max* | Soybean | 39 |
| *Gossypium raimondii* | Cotton | 27 |
| *Helianthus annuus* | Sunflower | 64 |
| *Hordeum vulgare* | Barley | 59 |
| *Leersia perrieri* | Cutgrass | 28 |
| *Lupinus angustifolius* | Blue Lupin | 24 |
| *Manihot esculenta* | Cassava | 26 |
| *Medicago truncatula* | Alfalfa | 29 |
| *Musa acuminata* | Banana | 33 |
| *Oryza sativa* (sp. Japonica) | Rice | 31 |
| *Panicum hallii* | Panicgrass | 28 |
| *Phaseolus vulgaris* | Green Bean | 18 |
| *Physcomitrium patens* | Club Moss | 14 |
| *Picea abies* | Norway spruce | 14 |
| *Populus trichocarpa* | Poplar | 25 |
| *Prunus persica* | Peach | 15 |
| *Selaginella moellendorffii* | Spike Moss | 16 |
| *Setaria italica* | Foxtail millet | 35 |
| *Solanum lycopersicum* | Tomato | 18 |
| *Solanum tuberosum* | Potato | 23 |
| *Sorghum bicolor* | Sorghum | 30 |
| *Theobroma cacao* | Cocoa | 16 |
| *Trifolium pratense* | Red Clover | 27 |
| *Triticum aestivum* | Common Wheat | 170 |
| *Triticum dicoccoides* | Emmer Wheat | 84 |
| *Triticum urartu* | Einkorn Wheat | 41 |
| *Vitis vinifera* | Grape | 33 |
| *Zea mays* | Corn | 37 |

### Characterization of the TaFMO family genes

The full wheat cDNA, CDS, and peptide sequences of candidate *TaFMO* genes identified were retrieved from IWGSC (The International Wheat Genome Sequencing Consortium (IWGSC) et al., 2018). Gene Ontology (GO) terms were retrieved from the Uniprot database (https://www.uniprot.org/; accessed January 2023). The structure of exons and introns were determined using the CDS and gene sequences of *TaFMO*s analyzed with the Gene Structure Display Server (GSDS) version 2.0 (gsds.cbi.pku.edu.cn; Hu et al., 2015). A small subset of putative *TaFMOs* were represented by splice variants and included for downstream analyses, on the basis that each variant could encode for fully functioning TaFMO proteins acting in different tissues, conditions, and developmental stages.

To assess whether the enzymatic activity of each candidate TaFMO could be carried out, the presence of three previously well-described FMO motifs (Eswaramoorthy et al., 2006; Mishina & Zeier, 2006; Schlaich, 2007; Thodberg & Jakobsen Neilson, 2020b) known to be crucial for cofactor binding and required for proper FMO function—the FAD- and NAD(P)H-binding motifs crucial for oxygenation activity, and the FMO-identifying motif facilitating in NAD(P)H-binding at the core pocket of the enzyme—were assessed with multiple sequence alignments (MSA) using MAFFT with the E-INS-I setting; see Appendix S2 (Katoh et al., 2019). Candidates that did not possess one or all motifs were retained in downstream phylogenetic analyses but flagged as ‘non-canonical’ (nc) FMOs with the assumption that oxygenation activity may be compromised, as we used conservation of all motifs as a proxy for predicting FMO function. We predicted the putative protein structure of all TaFMOs using the Protein Homology/Analogy Recognition Engine V 2.0 (Phyre^2^; Kelley et al., 2015) and then visualized them with PyMOL (Version 2.0 Schrödinger, LLC). We used the Simple Modular Architecture Research Tool (SMART; https://smart.embl.de/; Letunic et al., 2021) program to gather protein domains for display (Figure 2). Possible transmembrane domains and signal peptides for all TaFMO were predicted using the programs DeepTMHMM and SignalP - 6.0 (Hallgren et al., 2022; Teufel et al., 2022).

**Figure 2.**
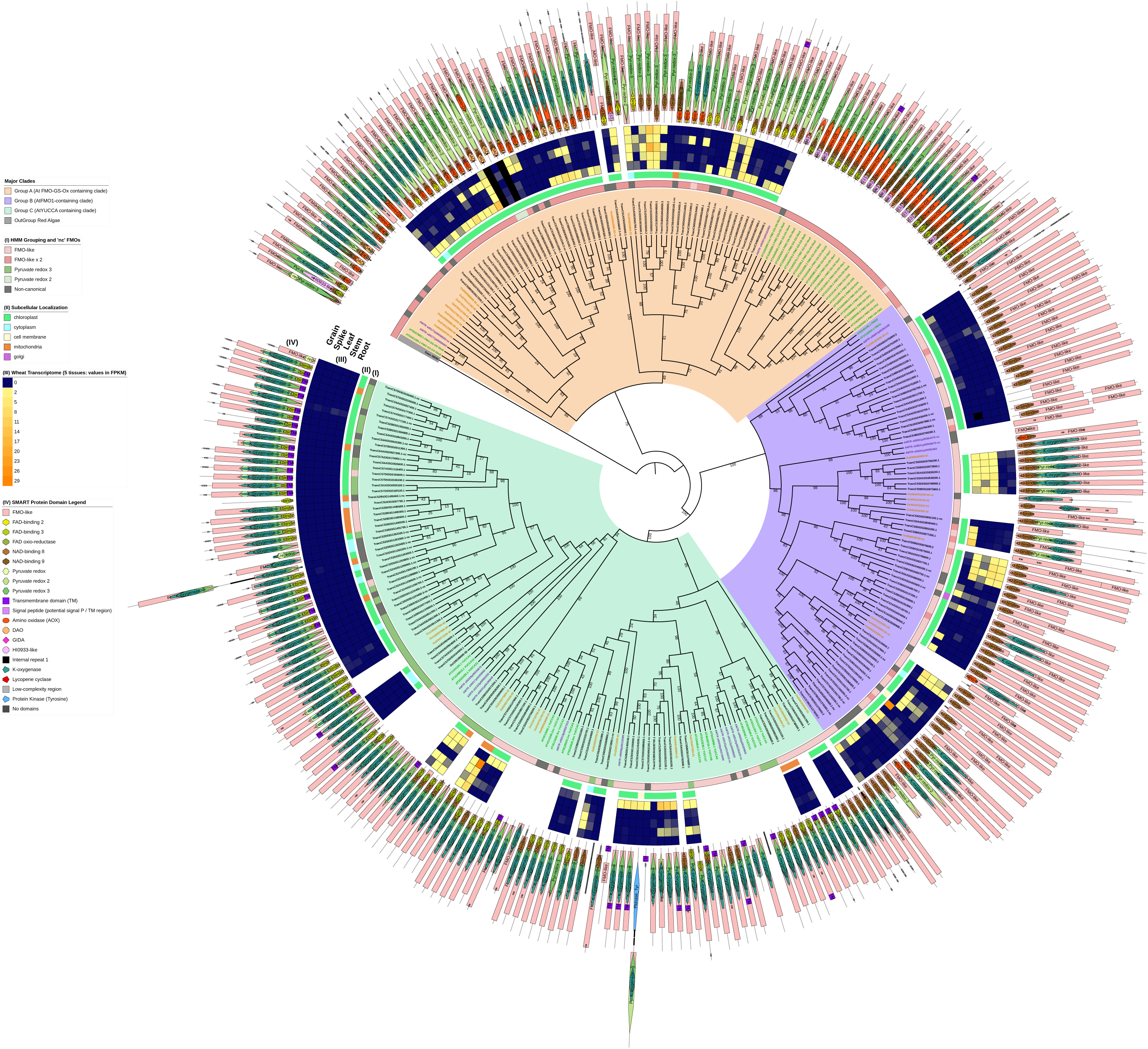
A maximum-likelihood (ML) consensus genealogy tree of total protein sequences of candidate TaFMOs (black gene IDs), OsjFMOs (yellow), AtFMOs (green), AmtriFMOs (purple), and GasuFMOs (grey). One GasuFMO (red microalgae) was used as outgroup. ML geneology was constructed from IQ-TREE (1000 standard bootstrap replicates, JTT+I+G4 substitution model). Three major subclades A (orange shading), B (purple) and C (green) were confidently supported across vascular plant FMOs. Outer circles: (I) shows the domain architecture the candidate FMO belongs to (from HMMER), as annotated in Figure 1; non-canonical ‘nc’ and truncated FMO protein candidates are marked with a dark grey box. (II) indicates predicted subcellular localization of TaFMO proteins. (III) represent transcript values (in FPKM) of each *TaFMO* candidate gene expressed under 5 tissue types (root, stem, leaves, spike, grain) from RNAseq experiments compiled on the Wheat Proteome database (Duncan et al., 2017). (IV) indicates SMART-predicted protein domains inclusive in each candidate FMO. Group A contains the least number of *TaFMO*s, totalling 55 splice variants pertaining to 39 genes (see S1 notes for explanation of exclusion of one *TaFMO* in the analysis), nine of which are categorized as ‘nc’. Group B contains a total of 63 splice variants pertaining to 57 genes, 14 of which are ‘nc’. Lastly, Group C contains 80 splice variants pertaining to 74 genes, 16 of which are ‘nc’.

### Phylogenetic analyses of the TaFMO gene family and sub-families

We inferred a maximum likelihood (ML) phylogeny to define relationships between total validated *TaFMOs* and validated *FMO*s from the model plant *A. thaliana*, the cereal crop *Oryza sativa* subsp. *japonica*, and sister group of all other angiosperms, *Amborella trichopoda*. We selected a red algae having a single *FMO* as an outgroup. We first aligned full-length FMO peptide sequences using MAFFT with the E-INS-i setting (Katoh et al., 2019). We validated the full-length alignment of the total FMOs across plant species using the software HOMO v2.0 (Jermiin, 2017) by using it to determine whether all FMO sequences met the phylogenetic assumption of evolution under reversible, stationary, and globally homogenous conditions (Jermiin, 2017; Jermiin et al., 2020). Following this, poorly aligned regions of the MSA were masked using the AliStat program (Wong et al., 2014) to ensure downstream analysis focused on regions where amino acid residues were readily alignable across sequences (see Appendix S2)(Wong et al., 2014). We then manually inspected alignments using Mesquite v 3.7 (Maddison and Maddison, 2023) and adjusted accordingly. Lastly, we used IQ-TREE v 2.2.0 to infer the ML tree based on amino acid alignments; we inferred the optimal substitution model using ModelFinder (JTT+I+G4), considering the Bayesian Information Criterion, BIC (Kalyaanamoorthy et al., 2017), and calculated branch support from 1000 standard bootstrapping replicates (Felsenstein, 1985).

We constructed separate ML phylogenies between all *TaFMO* and *FMO*s from closely-related species in the grass order, Poales—specifically, *H. vulgare* (barley), *T. urartu* (red einkorn wheat) and *O. sativa* ssp. japonica (rice) to improve our understanding of the relatedness of *TaFMO* in the three major clades A, B, and C (Figure S2-5). For detailed notes on alignment and modelling, see Appendix S2.

### Gene Duplication and Homeolog Analysis, and Protein Motif Analysis

We retrieved chromosomal locations of all *TaFMO*s in wheat from the RefSeqv2.1 gff3 file. We inferred homeologs based on well-supported phylogenetic clustering; separate sub-phylogenies were inferred for the three major partitions (clades A, B, and C) and a 70% bootstrap support value was used as a threshold to acknowledge clades with moderate support. Gene duplication and syntenic gene blocks of all *TaFMOs* (including ‘nc’ *TaFMOs*) were inferred through the MCSanX algorithm (Wang et al., 2012) via TBtools software v1.116 (Chen et al., 2022) and NOTUNG (Chen et al., 2000) to resolve tandem duplications and relationships between *TaFMO* homoeologous groups. Genes with no discernible relatives were denoted as ‘orphan’ *TaFMO* (Tables 2,3). A comprehensive synchronized map of all *TaFMO* homeologs (Figure 3) connected by designated group colours assigned in Figures S2-4 was generated using the shinyCircos software in RStudio (R Core Team, 2021; Yu et al., 2018). See Appendix S3 for more details. A *de novo* search for putative conserved protein motifs among all TaFMOs, HvFMOs, OsjFMOs, AtFMOs, AmTriFMOs, and PpFMOs was conducted with MEME Suite (Bailey et al., 2015): see Appendix S4.

**Figure 3.**
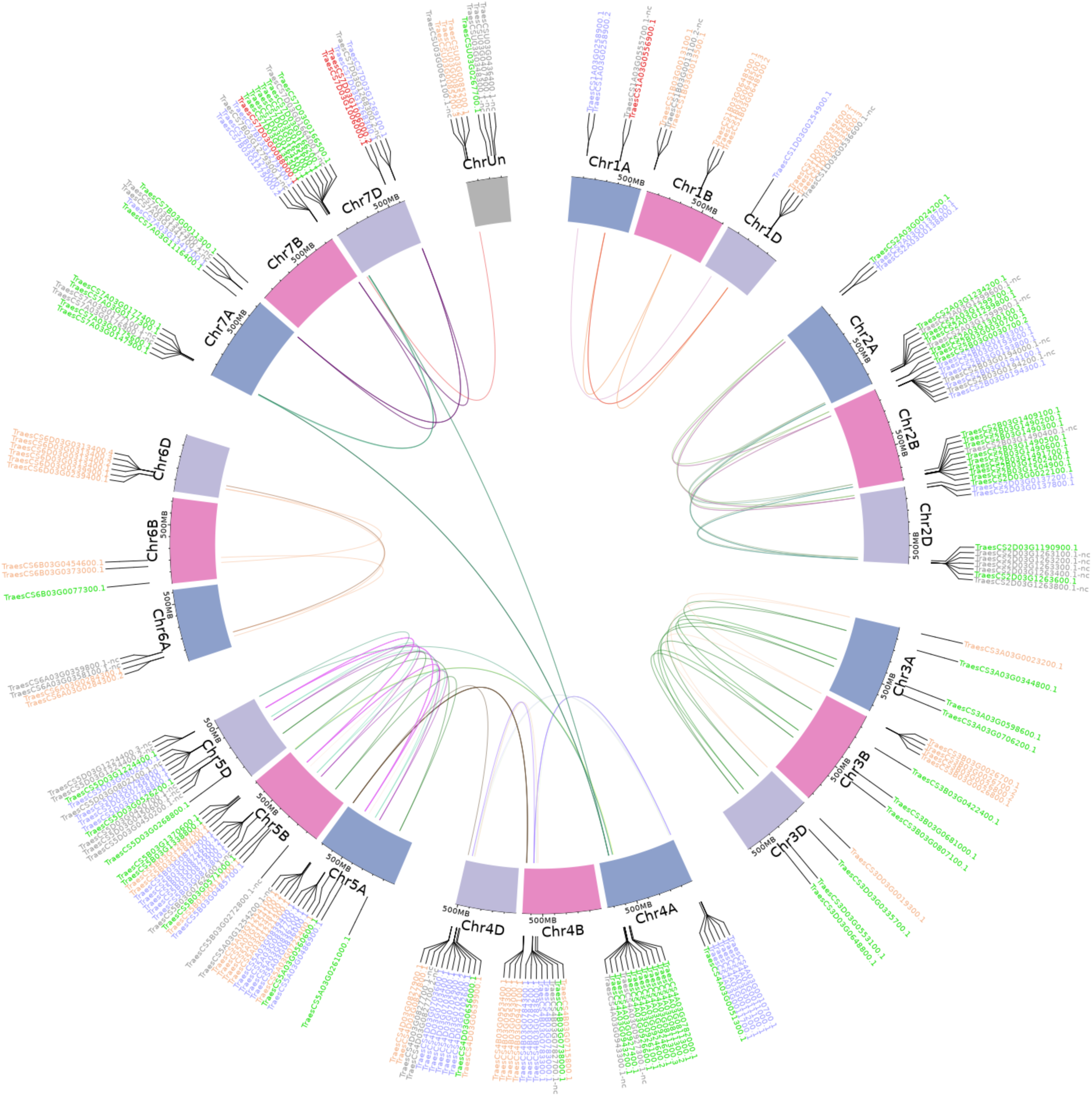
Genomic distribution of *TaFMO* groups A, B, and C candidates presented on a Circos plot (shinyCircos). Colours of gene labels correspond to the Group A (orange), B (blue) and C (green) partitions of *TaFMO* candidates. Red-labelled genes are orphaned/singleton *TaFMO*s, and grey-labelled genes are ‘nc’ *TaFMO*s. Cytobands exhibit chromosome tracks for chromosomes 1-7 (and unmapped chromosome, ChrUn) for each sub-genome A, B, and D in wheat. Links inside the circos plot are representative of whole-genome-duplication homeologs (syntenic triads) of wheat *TaFMO* genes delineated through a genome-wide synteny analysis; links are discretely coloured in correspondence to annotated subclades for each of the *TaFMO* candidates, seen in Figures S2-4 and S2 notes.

### TaFMO promoter regulatory elements analysis and transposable element identification

We searched promoter regions 1.5 kb upstream from the start codon of validated *TaFMOs* for plant *cis*-acting regulatory elements (PlantCARE: http://bioinformatics.psb.ugent.be/webtools/plantcare/html/; Lescot, 2002) and visualized putative cis-acting regulatory elements (CAREs) governing the different clades of *TaFMOs* (Tables S3,4, Appendix S5). We also compiled a list of putative transposable elements (TEs) detected by the CLARI-TE program (https://github.com/jdaron/CLARI-TE) either spanning introns and exons of *TaFMOs*, or flanking *TaFMOs* within a 2 kb region on either side (Table 4), from the TE annotation file provided in IWGSC RegSeqv2.1 (International Wheat Genome Sequencing Consortium et al., 2018; Zhu et al., 2021). The TE positions flanking or inside genes were transposed into *TaFMO* gene structure analysis. Refer to Appendix S6 for details.

### Transcriptome, proteome, and tissue localization data retrieval and display

Using public transcriptomic data across a range of conditions, we extracted candidate *TaFMO* gene expression information, obtaining transcript values in FPKM for five general tissues (root, stem, leaves, spike, and grain) and proteome data (peptide spectral counts) for different tissues from the developmental atlas in the Wheat Proteome database (Duncan et al., 2017). We acquired transcript data for a broad range of different conditions in values of log_2_ transcripts per million (tpm) for *TaFMO* expression under various abiotic stressors (cold, drought, heat) and biotic stressors (i.e., infection by *Septoria blotch, Fusarium graminearum,* or *Puccinia striiformis* f.sp. *tritici*) from the Wheat Expression Database (Ramírez-González et al., 2018) in order to survey the differential expression of *TaFMO* candidates, and infer their possible biological functions (see Tables S5-8 for an overview of studies and sources used).

## Results and Discussion

### FMOs are a largely expanded gene family in flowering plants

The *FMO* gene family is highly expanded in the vascular plants, as previously reported (Eswaramoorthy et al., 2006; Mishina & Zeier, 2006; Schlaich, 2007; Thodberg & Jakobsen Neilson, 2020a). Figure 1 exhibits the general number of potential FMO protein-encoding genes among select vascular plant reference genomes. The three most commonly present types of domain architectures that represent these plant FMOs are a single FMO-like domain (FMO-like), two consecutive FMO-like domains (FMO-like x 2), and a pyruvate redox 3 (Pyr3) domain. Some FMOs are predicted as fusion proteins, such as found in both wheat (*Ta*) and barley (*Hv*) with a tyrosine protein kinase conjugated to an FMO, alluding to a possible function in environmental sensing and downstream signalling coupled with oxidation processes (Shumayla & Upadhyay, 2023; Tichtinsky et al., 2003). The phenomenon of FMO fusion proteins does not present for lycophytes, moss, or algae, where total numbers of *FMOs* in the genome are substantially reduced relative to flowering vascular plant species. The total number of *FMOs* in plant genomes also appears to be largely reflective of species ploidy levels for recently diverged taxa, as exemplified in *Triticum urartu* (2n, 41 putative *TuFMOs*), *T. dicoccoides* (4n, 84 putative *TdFMO*s) and *T. aestivum* (6n, 170 putative *TaFMO*s). However, in plants considered diploid, the number of *FMOs* varies substantially, such as in *Arabidopsis* (29 *AtFMO*s), rice (31 *OsjFMO*s), and barley (59 *HvFMO*s), up to 64 potential *FMOs* in sunflower (*H. annuus*), which has double the amount of most diploid species in the eudicots (lower, green clade in Figure 1). We report several additional *FMO* candidates for rice compared to a recent report of 28 *OsjFMO*s by (Gaba et al., 2023), and for barley previously reported having 41 *HvFMO*s (Thodberg & Jakobsen Neilson, 2020b), highlighting the complexity of conducting genome-wide searches for highly expanded gene families.

An HMM search conducted against the RefSeqv2.1 wheat reference genome for cultivar ‘Chinese Spring’ yielded a total of 171 high confidence (HC) *TaFMO* genes, and 82 low confidence (LC) *TaFMO* genes. We selected 170 HC *TaFMO* genes including those deemed ‘unmapped’ to specific chromosomes (see Figure S1, Appendix S1 for additional information). A total of 30 TaFMO candidates are denoted as ‘non-canonical’ (nc), based on alterations in protein binding motifs previously reported as crucial for FMO function. The same logic was applied to sort for FMOs of *Arabidopsis thaliana* (*At*), *Oryza sativa* ssp. japonica (*Osj*)*, Hordeum vulgare* (*Hv*), and *Triticum urartu* (*Tu*).

### TaFMOs are consistently divided into three major groups across vascular plants, and span all chromosomes and sub-genomes

A maximum-likelihood phylogeny was constructed with protein sequences representing the total 170 *TaFMO* gene candidates (198 independent splice variants); *FMO*s from *Amborella trichopoda* (*Amtri*), rice *Oryza sativa* sp. Japonica (*Osj*), and *A. thaliana* (*At*) were included to probe for phylogenetic partitioning of *TaFMOs* among flowering plants. A red microalgae (*G. sulphuraria*) served as an outgroup, whose genome encodes for only one known *FMO* (Figure 1). We observed three major phylogenetic groupings, indicated as A, B, and C in Figure 2.

Domain architecture analysis (Figure 2, circle I) shows that Group A FMOs mostly share the same domain architecture type as the FMO from the outgroup red algae (FMO-like x 2), with two TaFMOs as exceptions having the pyruvate redox 2 architecture. Group B (purple) and Group C (green) FMOs form a clade across vascular plants with Group B mostly composed of an FMO-like architecture presenting as FMO-like x 2, whereas group C has a mix between the FMO-like architecture and pyruvate redox 3. The distribution of total FMOs across an array of vascular plants (or within one species) has been documented by Gaba et al. (2023), Thodberg & Jakobsen Neilson (2020b), and Yoshimoto et al. (2015), and the phylogenies presented in these studies also find Group B (*AtFMO1*-containing clade) and Group C (‘*YUCCA*’ clade) to be more closely related to each other than to Group A (the *AtFMO GSOX-*containing clade).

We found that *TaFMOs* span all seven (and unmapped) chromosomes and sub-genomes (A, B, and D) derived from progenitor grass species *Triticum urartu*, *Aegilops speltoides*, and *Aegilops tauschii*, respectively (Ramírez-González et al., 2018; T. Zhu et al., 2021). Chromosomes 2, 4 and 7 are enriched with *TaFMO*s in the telomeric regions (Figure 3). Translocations of *TaFMOs* are inferred to have occurred most frequently between chromosomes 4A, 7A, and 7D, which are reported to be hot spots for homoeologous gene translocation events for many genes in wheat (Zhou et al., 2020)(Appendix S3).

### Sub-groups reveal non-uniform patterns of TaFMO gene expansion

Tables 2 and 3 summarize the synteny status of Groups A, B, and C *TaFMO* homeologs and homeolog sub-genome ratios (denoted as a *TaFMO* ratio of 1 : 1 : 1 from A, B, and D sub-genomes), respectively. Group A *TaFMOs* show substantially less syntenic triads with only 8.2% of triads in Group A exhibiting an even ratio of 1 : 1 : 1; even more, around 42% of triads in Group A were missing either an A-, B-, or D-copy, a higher frequency than the 13.2% expected for all wheat proteins in the genome (Table 3). Group C, subclade C1, showed the most conserved ‘clean’ homoeologous triads (56.3%) such that there were rarely any singleton/orphan or ‘nc’ *TaFMOs*, and each *TaFMO* triad had corresponding orthologues from other grass species (Figure S4); the high evolutionary conservation of these *TaFMO* triads is perhaps due to functional constraints influencing gene retention. In contrast, *TaFMOs* in subclade C2 (along with barley *FMO*s) appear to have large, uneven homeolog expansions via tandem duplications and intra- or interchromosomal dispersion of genes (Figure S4).

**Table 2.** *TaFMO* Homeolog analysis and literature evidence. *[is presented in the supplementary excel file]*

**Table 3.**
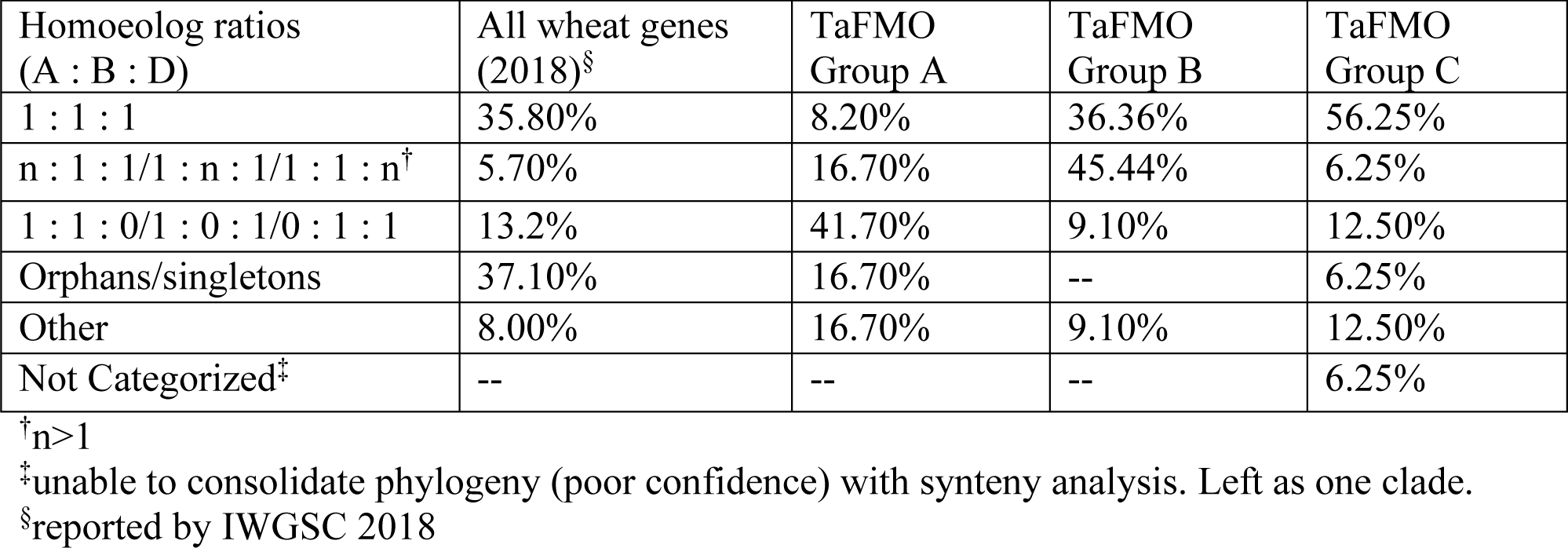
*TaFMO* Homeolog (triad) analysis.

In Group B, 36.4% of *TaFMOs* belong to even triads which more closely resemble the anticipated percentage (35.8%) of all predicted wheat proteins belonging to even homoeologous triads (Table 3)(Ramírez-González et al., 2018). Group B appears to have many instances of sub-genome homeolog expansion and uneven tandem duplications (Table 2). Subclade B1 appears to contain three pairs of *TaFMO* WGD homeologs that underwent multiple duplications, with more *HvFMO* orthologues present than *OsjFMO* and *TuFMO*, (Figure S3), while subclade B2 shows *FMO* expansions for all grass species surveyed, but none for *Arabidopsis* (Figure S3).

Such divergent patterns of gene retention and expansion among homoeologous triads between the three major groups of *TaFMOs* might implicate differing biological roles. (Birchler & Yang, 2022) make a point that genes involved in defense mechanisms are more likely to diverge after duplication events; the retention of the active sites for some of the highly duplicated Group B triads could infer a role in defense-related functions.

### Variations in canonical motifs delineate the FMOs in wheat and related grasses

We detected fifteen of the most conserved protein motifs in FMOs across *Arabidopsis* and the grasses rice (*Osj*), barley (*Hv*), red einkorn (*Tu*) and wheat (Figure S5). FMOs in plants are reported to contain three crucial protein motifs essential for ‘canonical’ FMO activity; these are the FAD-binding motif (GxGxxG), the NAD(P)H-binding motif (GxGxxG), and the FMO-identifying motif (FxGxxxHxxxY/F; (Eswaramoorthy et al., 2006; Hansen et al., 2007; Huijbers et al., 2014)). While the FAD- and NAD(P)H-binding sites are active sites surrounding the enzyme pocket where hydroxylation takes place, the FMO-identifying motif is considered a linker region between the two active sites (Malito et al., 2004). The last and less widely talked about motif is the ‘F/LATGY’ motif, more recently referred to in some studies as the ‘ATG-containing’ or ‘TGY’ motif (Gaba et al., 2023; Yoshimoto & Saito, 2015); this motif is thought to be a crucial factor governing *N*-hydroxylation (Fennema et al., 2016; Stehr et al., 1998).

The FAD-binding motif at the N-terminus is very well conserved with regards to three glycine residues across all groups (Figure 4a,b). However, the residues surrounding this motif for Group A and subclade B2 from Group B is recorded to be distinct from that in subclade B1 and Group C (Figure 4a,b; Figures S2-4); the FAD-binding site variant 1 (for Group A and B2) and the other variant are highlighted in orange in Figure 4a,b. An extra copy of the FAD-binding variant 1 also exists between the NAD(P)H-binding site and the ATG-motif in most of the examined grass species (*Hv, Tu*, *Ta*) in subclade C2, but not for rice. This may indicate a novel function for FMOs specific to Triticeae. The NAD(P)H-binding motif across all FMOs (Figure S5) is well-conserved for the first glycine residue, but not for the other two glycine residues (Gxgxxg; Figure 4a). The FMO-identifying motif for FMOs in Group A has the first residue swapped from the canonical F (Phenylalanine) for W (Tryptophan), which is the case for most of the plant species examined (Figure 4a). Additionally, the ATG-containing motif for Group A FMOs across all flowering plants surveyed are highly conserved, represented as HCTGY or YCTGY. The differences of the Group A ATG-motif and FMO-identifying motifs relative to Groups B and C may allude to specificity for *S*-oxygenation of compounds and specialized metabolites yet uncharacterized in cereals, where Group A FMOs from *A. thaliana* (FMO-GSOX enzymes) are well-characterized for their *S*-oxygenation activity during synthesis of specialized metabolites (Hansen et al., 2007). The ATG-containing motif is most varied for Group B FMOs, where in addition to the canonical ‘(F/L)ATGY’ residues reported by (Schlaich, 2007), other variations are present (LATGF, FLATGF, FATGY, LATGY, FGTGF) in the B2 expanded subclade comprising other grass species (*Tu, Hv, Osj*). By contrast, Group C FMOs mostly present with ‘LATGY’ motifs (and some ‘MATGY’ motifs in barley and wheat) in subclade C1 and ‘FATGY’ motifs in subclade C2 (Figure 4a).

**Figure 4.**
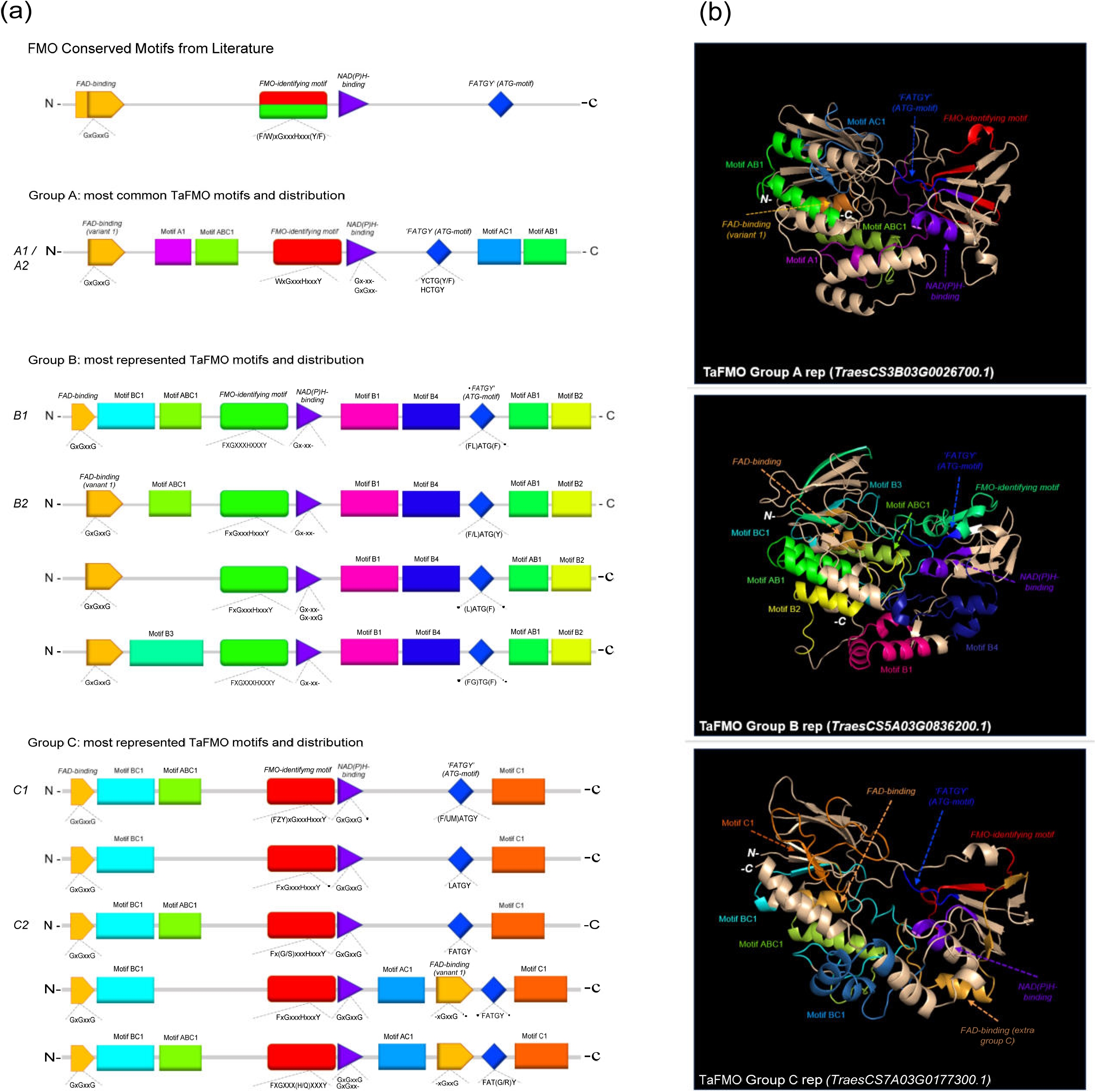
(a) Consensus MEME motifs for conserved and unique motifs across Groups A, B, and C TaFMO proteins. (b) Predicted 3D protein structures for Groups A, B, and C TaFMO with highlighted motifs.

Beyond the documented protein motifs discussed above, one novel motif is reported for each Group A and Group C FMOs (referred to here as motif-A1 and motif-C1, respectively; see Figure S5 for these and other motifs referred to in this section). Meanwhile, Group B FMOs have four clade-specific motifs (motif-B1-4) (Figure 4). Groups A, B, and C FMOs all share one novel motif each with each other (motif-AB1, motif-AC1, motif-BC1). Additionally, there is a novel motif common to almost all FMOs located at the N-terminus following the FAD-binding site for most FMOs, reported here as motif-ABC1 (Figure 4a). A look at predicted protein models highlights that many of these novel motifs may present as-yet-unclassified linker regions contributing to the integrity of the enzyme, or motifs that govern substrate specificity and binding affinity (Figure 4b). We hypothesize that differences in tertiary protein structures, including these clade-specific motifs, likely play a crucial role in substrate specificity (Li et al., 2008; Zhu et al., 2019). While no plant FMOs have been crystalized to date, the use of 3D protein homology modelling and ligand-docking simulations—in corroboration with functional analyses—may shed some light on which compounds TaFMO, and other cereal FMOs, may be acting upon.

### Possible roles of Group A and C TaFMO: from pathogen defense to grain development

Group A *FMO*s contain all the *Arabidopsis FMO-GSOX* (glucosinolate oxidase) genes. These FMOs have been experimentally validated to participate in glucosinolate (GSL) biosynthesis by catalyzing the *S*-hydroxylation of methylthioalkyls to methylsulfinylalkyls, where GSLs are a class of specialized sulphur-containing metabolites involved in plant-herbivory defenses (Cang et al., 2018; Hansen et al., 2007; Kong et al., 2016; Li et al., 2011; Schall et al., 2020). GSLs are predominantly found in the mustard seed family (Brassicaceae), although there is evidence that 500 neighbouring eudicot species outside this family may also contain one or more of the 120 documented GSLs (Flamini, 2012). The possible metabolites that these Group A FMOs help modify in cereals are not well-understood, as cereals are not reported to produce GSLs (Hansen et al., 2007; Thodberg & Jakobsen Neilson, 2020a). More recently, other *A. thaliana* FMOs have been documented to synthesize trimethylamine *N*-oxide (TMAO)(Fennema et al., 2016), the levels of which are increased in plants under low temperatures, drought, and high salt stress conditions (Catalá et al., 2021). Group A *FMO*s segregate into two major subclades, A1 and A2 (Figures 2, S2). While little is known for Group A *FMOs* in barley and red einkorn, it is reported that five rice *FMO*s from subclade A1 (*Os01t0368000-nc, Os02t0580600-nc, Os07t0111700, Os07t0111900, Os07t0112000*) are co-expressed with *WRKY13* (a transcription factor involved in pathogen defense) upon attack by rice sheath-infecting fungi (John Lilly & Subramanian, 2019). For *TaFMOs* in subclade A1, the literature supports involvement in root drought tolerance, tiller growth, and nutrient accumulation in the grain (Table 2, Figure S6). The rice *FMO Os10t0553800* in subclade A2 may play an important role in rice germination and seedling establishment during floods, via epigenetic methylation responses to anaerobic conditions (Castano-Duque et al., 2021). By contrast, several *TaFMOs* in subclade A2 are implicated in disease resistance against both biotrophic (Navathe et al., 2022) and necrotrophic pathogens (Nussbaumer et al., 2015), in addition to root drought tolerance (Figure S6). The amino oxidase (AOX) protein domain which is seen in almost all FMOs in Group A at the N-terminus but rarely for Group B (and which is missing from Group C), is larger in subclade A2 AtFMOs (∼150 amino acid residues), but not for wheat (∼60 residues) or other plant FMOs (Figure 2). The AOX domain in Group A TaFMOs appears to encompass the motif-A1 mentioned previously unique to Group A and may participate in substrate binding specificity for novel compounds in wheat.

Group C contains the “*YUCCA*” (*YUC*) genes that are known to participate in auxin (idole-3-acetic acid, IAA) biosynthesis (Cao et al., 2019; Kendrew, 2001a), where auxin is involved in many plant development processes (Chandler, 2015; Schlaich, 2007). We identified 74 *TaFMOs* in Group C, which can be split into two main subclades, C1 and C2. Several *TaFMOs* in subclade C1 are implicated in the response to drought stress (Table 2, Figure S7), found to be upregulated during root drought treatment (Grzesiak et al., 2019), or associated with an increased reactive oxygen species (ROS, H_2_O_2_) burst in response to root drought conditions (Kamruzzaman et al., 2022). In subclade C2, several *TaFMOs* are implicated in plant development, including in grain (Mangini et al., 2021) and seed development (Li et al., 2014), and tiller number increase (Yu et al., 2021). To date, only three homoeologous genes involved in seed development have been functionally characterized via gene cloning (Li et al., 2014). A catalogue of 63 ‘*TaYUC’* gene candidates involved in auxin biosynthesis by Yang *et al*. (2021) included several *TaFMO* genes outside Group C (dispersed throughout subclade B2; Table S4), hinting towards a flexibility in *TaFMO* involvement in auxin biosynthesis-regulated plant development. In the absence of a full gene family survey and experimental characterization, adopting a more wheat-specific nomenclature for the *FMO* gene family via the guidelines put forward by the Wheat Initiative community (Boden et al., 2023), may minimize confusion on gene names in wheat to streamline future research efforts.

### Group B TaFMOs: major players in broad-spectrum disease resistance

Group B *FMO*s can be split into two distinct subclades, B1 and B2 (Figure 2). This group contains the *Arabidopsis AtFMO1* and *AtFMO2* genes. AtFMO1 hydroxylates the N atom of an L-lysine catabolic product to form *N*-hydroxylated pipecolic acid (NHP), crucial for establishing systemic acquired resistance (SAR) to combat microbial pathogen invasions (Chen et al., 2018; Conrath, 2006; Hartmann et al., 2018; Mishina & Zeier, 2006), and a lack of NHP (due to defects in its biosynthesis) results in immunocompromised *Arabidopsis* plants. Various studies have recently contributed to better understanding the mechanism of NHP-mediated SAR regulation in other plants, including cereal crops, by searching for functional orthologs to *AtFMO1* and their possible involvement in disease resistance (Holmes et al., 2019; Lenk et al., 2019; Schnake et al., 2020; Vlot et al., 2020; Zhang et al., 2021).

In subclade B1*, AtFMO1* and *AtFMO2*, the only Group B *FMOs* in *Arabidopsis,* form a clade with 18 *TaFMOs*, where only three candidates are ‘nc’ (Table 2, Figure 4b). All FMOs from subclade B1 possess the most minimal number of known protein domains via SMART analysis, whereas multiple protein domains are present for TaFMO from all other clades (Figure 2). Of note, AtFMO1 harbors an additional K-oxygenase domain (IPR025700) in the first half of its protein sequence, which is not seen in any of the B1 TaFMOs or AtFMO2 but is present in most B2 TaFMOs and TaFMOs in Groups A and C. The K-oxygenase domain is involved in siderophore biosynthesis, and in ornithine and lysine hydroxylation (Krithika et al., 2006; Visca et al., 1994); therefore, presence of this K-oxygenase may play a critical role in hydroxylation of L-lysine and/or L-ornithine derived compounds.

While subclade B1 *TaFMOs* show little to no transcriptional activity in the conditions surveyed, certain subclade B2 *TaFMO*s exhibit higher transcriptional activity (Figure 2, Figure 5 – III/IV). The *TaFMO TraesCS5D03G0799800.1* in subclade B2 (B2β-3b) was detected as the most transcriptionally upregulated “FMO1-like” *TaFMO* in response to treatment of wheat coleoptiles with NHP (Zhang et al., 2021). Two homoeologous triads in B2α-2 (Table 2) show transcriptional upregulation in root, stem, and leaf tissues (Figure 2), and peptide abundance of these triads was also detected in a wide variety of wheat tissues (Figure 5 – II). These *TaFMOs* in B2α-2 undergo transcriptional activation in wheat seedlings during infection by the fungal pathogens *Zymoseptoria tritici* (*Zt*)(Rudd et al., 2015; Yang et al., 2013) and *Puccinia striiformis* f.sp. *tritici* (*Pst*)(Cantu et al., 2013; Dobon et al., 2016), but have very little to no transcriptional activation during older developmental stages of wheat, such as seen in controls during infections by *Fusarium graminearum* (*Fg*) in the wheat grain head (Schweiger et al., 2016) and *Magnaporthe oryza* (*Mo*) at the grain-filling stage (Islam et al., 2016) (Figure 5 - III).

**Figure 5.**
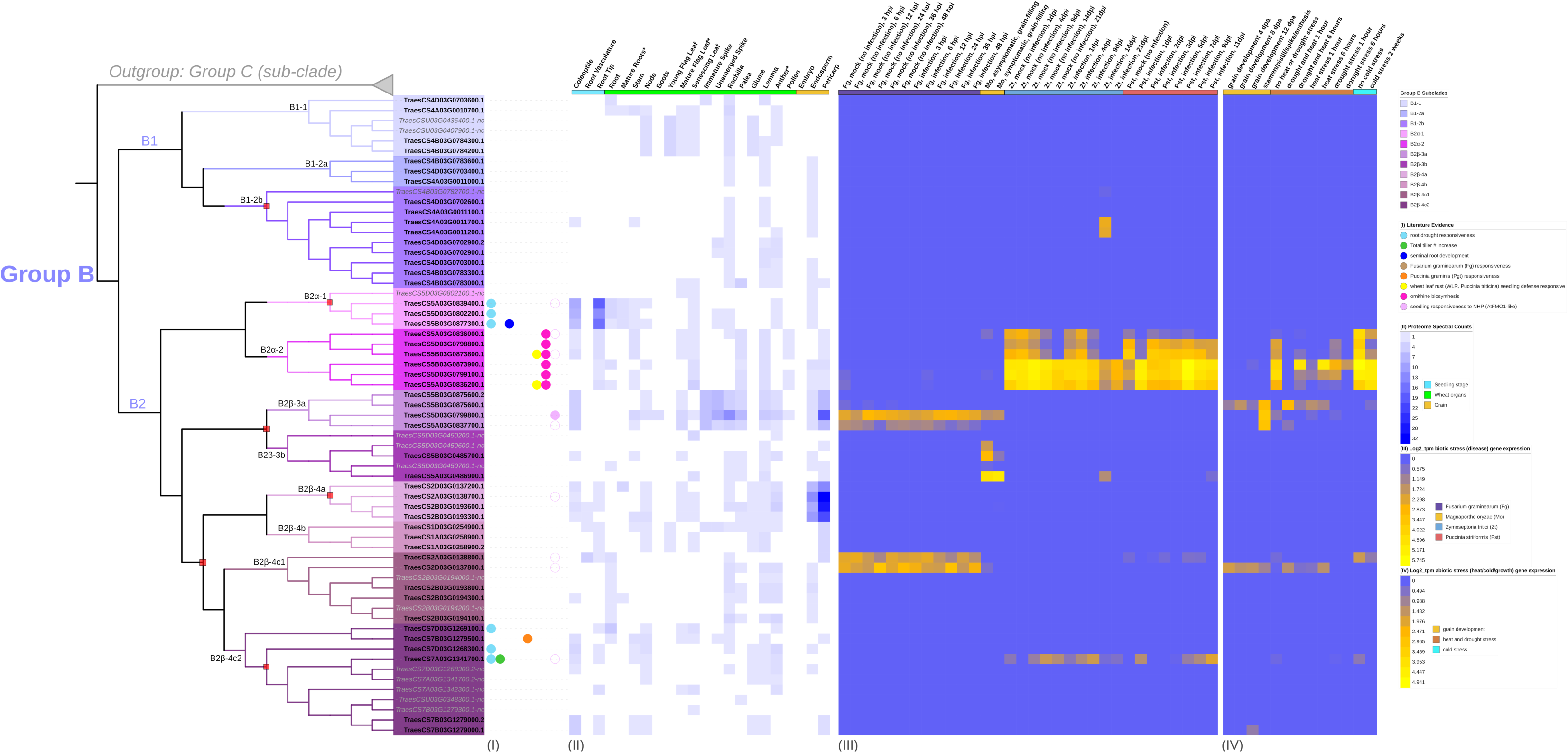
Transcription of group B *TaFMOs* under various conditions explored from literature data. (Columns I) reports of previous evidence of transcript detection or cloning under various conditions described in the legend. (Columns II) Protein expression (spectral peptide counts) for each candidate in various tissues from https://wheatproteome.org/ (Duncan et al. 2017). (Columns III) Heatmap of *TaFMO* transcripts in response to four types of biotic stresses; response to *Fusarium graminearum* infection(*Fg*), mock inoculation versus pathogen infection in a time-course of 3, 6, 12, 24, 36, and 48 hours post infection (hpi); *Magnaporthe oryza* (*Mo*), asymptomatic (control) versus symptomatic (infection) conditions at grain-filling stage; *Zymoseptoria tritici* (*Zt*), mock inoculation versus pathogen infection in a time-course of 1, 4, 9, 14, 21 days post infection (dpi); *Puccinia striiformis* f.sp. *tritici* (*Pst*), uninfected control versus pathogen infection in a time-course of 1, 2, 3, 5, 7, 9, 11 dpi. (Columns IV) Heatmap of *TaFMO* transcript response to grain development over a time-course (4, 8 12 days post anthesis), heat and drought stress (1 and 6 hours), and cold stress (control versus 2-week cold stress). Heatmap data is displayed as Log2 transcripts per million (tpm), with blue being 0 and higher-fold expression approaching bright yellow (see legends).

One of the B2α-2 homoeologous triads (*TraesCS5A03G0836200.1, TraesCS5B03G0873900.1, TraesCS5D03G0799100.1*) shows greater upregulation of transcription after *Pst* infection at one day post-infection (dpi), which is sustained through 11 dpi; *TraesCS5A03G0836200.1* is also reported by (Vikas et al., 2022) in a genome-wide association study (GWAS) to be likely involved in seedling resistance to the biotrophic fungal pathogen *Puccinia triticina* (*Pt*), the wheat leaf rust pathogen closely related to *Pst*. Another *TaFMO* in sub-clade B2β-4c(2), *TraesCS7B03G1279500.1*, might be involved in *Puccinia graminis* f.sp *tritici* (*Pgt*, stem rust) resistance (Sahu et al., 2021). In a QTL analysis by (Matros et al., 2016), the two triads in subclade B2α-2 were also detected as significantly enriched in SNPs associated with ornithine metabolism in wheat. Ornithine is an amino acid bearing great resemblance to lysine and has functional significance in plant abiotic stress tolerance (Kalamaki et al., 2009) through its role in osmoregulation during drought and salinity stress (Roosens et al., 1998; Xue et al., 2009). These two triads are among the only Group B *TaFMOs* to be transcriptionally responsive in the abiotic conditions presented, downregulated during drought stress, heat stress, and combined drought + heat stress at one hour, but bounce back to nearly control levels after six hours of treatment for certain splice variants (Figure 5 – IV)(Liu et al., 2015). Transcripts of the *TaFMO* triad in B2α-1 (except for the ‘nc’ 5D homeolog) have been detected in wheat roots (Figure 2, Table 2). (Grzesiak et al., 2019) reported that genes of this triad were upregulated in response to water deprivation in wheat root tissues, though no upregulation of transcription in leaf tissues was detected in the drought study by Liu et al. (2015) (Figure 5 – IV), highlighting a need for careful interpretation of tissue-specific expression patterns of *TaFMOs*. Furthermore, a GWAS analysis indicates that the B-copy gene of this triad, *TraesCS5B03G0877300.1*, influences root-growth and biomass yield-related traits (Zhao et al., 2023). Trends for other candidates of interest are found in Appendix S7.

In summary, several lines of evidence hint at roles in responsiveness to various microbial diseases for several *TaFMOs* in subclade B2. Additionally, certain subclade B2 *TaFMOs* may operate during plant development (tiller increase) and drought, heat, and salinity tolerance. Thodberg et al. (2020) showed that a novel Group B FMO in fern (FOS1) participates in both the *N*-hydroxylation of a novel class of metabolites not previously reported in vascular plants and in the biosynthesis of cyanogenic glycoside. Likewise, the exploration and characterisation of these B2 *TaFMOs* may identify novel functions and metabolites towards plant development, biotic, and abiotic stressors.

### TaFMO UTRs are populated with elements involved in complex regulation

Untranslated regions in transcripts (UTRs) have been reported in plants as important regulatory regions often harbouring important *cis*-acting regulatory elements (CAREs) that influence gene expression, modulate translational efficiency, and increase the coding capacity of genes that have different splice variations (Mignone et al., 2002; Srivastava et al., 2018). We found a wide variety of CAREs in the promoter regions 1.5 kb upstream of the start codon of *TaFMO* genes (Figures 6, S8, S9). The presence and sizes of 5’- and 3’-UTRs among *TaFMOs* varied; we observed that for all *TaFMOs,* the 3’-UTRs were longer than the 5’-UTRs, when UTRs were present (Figures 7, S10, S11). Srivastava *et al*. (2018) reported elongated UTRs (particularly 3’- UTRs) in rice compared to *Arabidopsis*. In wheat, 3’-UTRs have also been reported to have a critical involvement in mediating drought stress (Ma et al., 2023) and in establishing resistance against stripe rust (Zhang et al., 2019), possibly through regulating mRNA stability. Of note, all the tandemly duplicated wheat *TaFMOs* in subclade C2 lack any UTRs (Figure S11) and show virtually no (or low) transcript responsiveness to the biotic and abiotic conditions explored (Figure S7 – III, IV), with minimal presence of expressed peptides detected in various tissue types.

**Figure 6.**
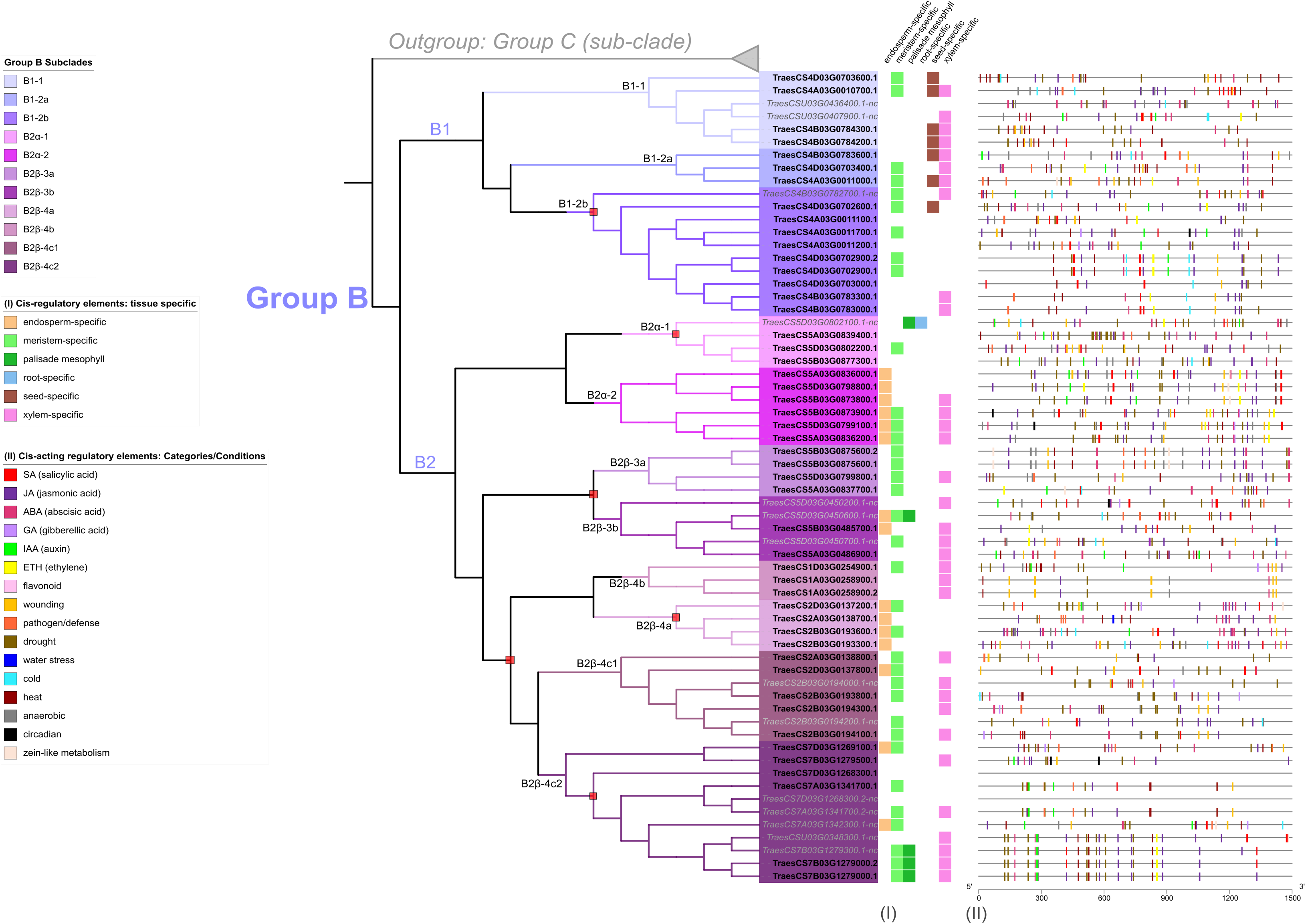
Plant *cis*-acting regulatory elements (CARE) detected in a 1.5 kb region upstream of start codons for all *TaFMO* in Group B. X-axis denote the number of base pairs away from the start codon, where 5’ is 1,500 bp upstream ATG start codon (3’ and 0 bp is right before the ATG start codon). Polytomy regions are denoted by a red box. Presence of regulatory elements specific to different tissues are described in (I). Other responsive promoter elements are denoted in the figure legend and displayed as rectangles of corresponding colour to the condition in (II).

**Figure 7.**
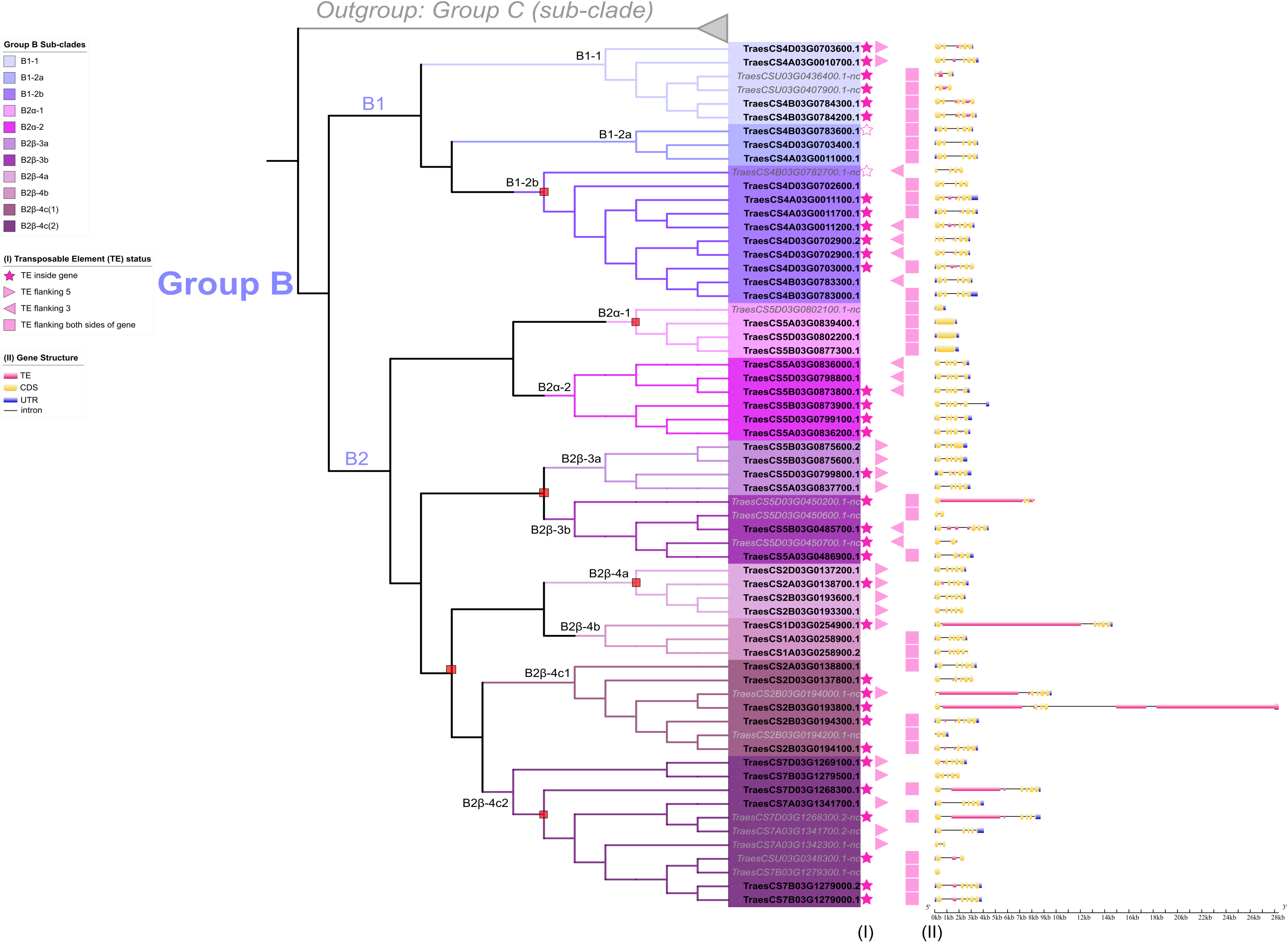
Graphical display of the gene structure and transposable element analysis for *TaFMOs* in Group B. Polytomies are denoted by a red box at respective nodes on the ML tree. Status of TE distribution is described in (I) for filled star: TE insertion inside gene; unfilled star: non-coding region of TE spanning entire gene; filled triangle: TE flanking either 5’ or 3’ UTR of gene; and filled square: TE flanking both sides of the gene. A 2 kb region upstream of the start codon and downstream of the stop codon were scanned for flanking TEs. (II) The gene structure is displayed for each *TaFMO,* with graphical depiction of TE insertion, if present. The X-axis scale describes the size in kb of nucleotides for each gene region. In Group B, exon numbers vary from three to five (B1), one (B2α-1), and one to five (all rest of sub-clade B2).

Focusing on the Group B *TaFMO*s, analysis of the CAREs revealed involvement in responsiveness to major phytohormones and conditions (Figure 6, Appendix S5). Jasmonic acid (JA)-responsive CAREs were present in all *TaFMO* promoters at one or multiple sites; JA is a phytohormone known for involvement in many plant processes ranging broadly from photosynthesis to heat stress-responsiveness (Per et al., 2018), but most widely documented for defense against necrotrophic pathogens (Qi et al., 2016) and induced systemic resistance (Heil, 2002). In wheat, JA-signalling has also been implicated in seedling salt-stress tolerance (Qiu et al., 2014). Other notable phytohormone-responsive CAREs in Group B are for salicylic acid (SA), gibberellic acid (GA) and ethylene (ETH) (Tables S3,4, Figure 6). SA is a major phytohormone involved in SAR and pathogen defense signalling pathways (Dey et al., 2014; Ding et al., 2018). SA-responsive CAREs were found in higher abundance and closer to the start codons of the triads from B2α-2 (Figure 6, Table S4), whose genes were more transcriptionally active in *Zt-* and *Pst*-infected wheat (Figure 5 – III); this may indicate that number and location of these CAREs influence which *TaFMOs* are activated under pathogen stress. GA is a phytohormone that was documented in wheat for its involvement in enhancing disease resistance against *F. graminearum* infection, by modulating both primary and specialized metabolism during early plant defense signalling (Buhrow et al., 2021); GA-responsive CAREs are found predominantly in subclade B2, further supporting that *TaFMOs* in this subclade are likely involved in microbial disease resistance. Auxin (IAA)-responsive CAREs are found to be abundant among many Group B *TaFMOs* (Table S4), challenging the idea of Group C *TaFMOs*—the ‘*YUCCA*’ *FMO* clade—being largely responsible for auxin biosynthesis and growth regulation in plants for underreported crop species (Kendrew, 2001b).

### Transposable elements, gene structure diversity, and differential TaFMO expression

We explored transposable element (TE) distribution in or near the identified *TaFMO* genes and affecting intron-exon structure to find support for relationships among *TaFMO* family members (Figure 7, Figures S10,11; Appendix S6). Focusing again on the Group B *TaFMOs*, we observed much variation in the number and spatial distribution of introns and exons; high similarity of gene structures was found among certain syntenic triads highlighting distinct *TaFMO* subfamilies that possibly remain conserved for functional importance. In contrast, extensive intron-exon shuffling following gene duplication events (França et al., 2012; Kolkman & Stemmer, 2001) may indicate lower selective pressure and neofunctionalization, such as in the case for subclades B2β-4c1 and B2β-4c2 (Figure 7).

Nearly all *TaFMO*s (90%) are flanked by TEs within a 2 kb region upstream and/or downstream of the start or stop codons (Figure 7, Table 4) in accordance with a report of TEs representing over 80% of the wheat genome (Bariah et al., 2020). 44.7% of all *TaFMO* genes show disruption by TE insertion, mostly occurring inside introns, inside 5’- or 3’-UTRs, or, very rarely, inside exons (Figure 7). TE insertions are present in 56.1% of Group B *TaFMO*, with the largest TE interruption in *TraesCS2B03G0193800* increasing its total gene length to upwards of 29 kb (Figure 7). Most TE disruptions in subclade B2 occur in the B- and D-copy homeologs and appear to be correlated with differential expression patterns among homoeologous triads. Varying locations of TE insertions in the highly tandemly duplicated B-copy *TaFMO*s in B2β-4c(1) correlate with no transcriptional expression compared to the A- and D-copies, and only an A-copy *TaFMO* in B2β-4c(2) lacking TE insertions was transcriptionally active (Figure 5 – III, IV; Figure 7). In contrast, one of the triads in B2α-2 (*TraesCS5A03G0836200, TraesCS5B03G0873900, TraesCS5D03G0799100*) shares the same TE insertion in its three members, which are all more transcriptionally active than gene members in the other B2α-2 triad lacking this TE insertion, particularly upregulated towards *Pst* infection and cold stress responses (Figure 5 – III, IV; Figure 7).

**Table 4.** Transposable element status for *TaFMO* in Group A, B, and C.

| **Table 4. Transposable element status for *TaFMO* in Group A, B, and C** |  |  |
| --- | --- | --- |
| FMOs in *T. aestivum* cv. Chinese Spring (RefSeqv2.1) | TE insertion<br>inside | TE flanking (5' UTR, 3' UTR,<br>both) |
| Total *TaFMO* (170) | 44.7% | 90.0% |
| -- |  |  |
| Group A TaFMO | 60.0% | 100.0% |
| Group B TaFMO | 56.1% | 91.2% |
| Group C TaFMO | 27.0% | 85.1% |

Yang *et al*. (2021) reported on the gene structure of a candidate ‘*YUCCA*’ *FMO* in wheat, *TaYUCCA5-D* (*TraesCS1D03G0254900.1*, Group B), containing a first intron size of approximately 13 kb. We now know this is due to a TE insertion (Figure 7), underscoring a need for expanding on factors driving gene structure diversity in large gene-family analyses. We see that TE insertions in the intronic and 3’-UTR regions may play a role in influencing gene expression of triads on a sub-genome level, possibly leading to eventual neofunctionalization of *FMO* family members. Several studies indicate that TEs are important drivers in polyploid plant evolution (Vicient & Casacuberta, 2017; Gill et al., 2021), such as in wheat where distinct TE families are correlated with up- or down-regulation of genes in their vicinity (Bariah et al., 2023). Thus, exploring classes of TE elements that lie in or around *TaFMOs* could aid in predicting gene expression regulation in the different wheat sub-genomes.

## Concluding Remarks

We present here a comprehensive phylogenetic analysis of the *FMO* superfamily in wheat and integrate various publicly available transcriptomic and proteomic data to shed light on the breadth of expansion for this superfamily in wheat and predict potential roles for the members of this family for hypothesis generation and further functional studies. We show that *TaFMOs* segregate into three distinct groups, and that unique domain architectures and protein motifs may indicate ligand-binding specificities among sub-groups of *TaFMOs*. We corroborate the expression data from existing literature of certain candidate *TaFMOs* with their possible biological functions as discussed in various studies. This study provides a solid foundation for the further exploration and functional characterization of the *FMO* gene family in common wheat.

## Data Availability

The raw FASTA files, multiple sequence alignments (MSA), phylogenetic tree files, HMM search/scan output files, and SMART protein domain files are available upon request.

## Acknowledgements

We thank our funding support by the Natural Sciences and Engineering Research Council of Canada (NSERC) CREATE program, as well as the NSERC CGS-D program. The authors are indebted to Dr. Sean Graham for help reviewing the phylogenetic analyses, manuscript editing and valuable discussions. Additionally, we thank Dr. Matthias Kretschmer for his constructive feedback.

## Supporting Information (Figures and Appendix Notes in SI PDF file and Tables in Excel file)

**Appendix S1.** Sources for sequence data and additional information on gene differences in *TaFMO* between IWGSC RefSeqv1.2 and RefSeqv2.1, and metadata.

**Appendix S2.** Alignment and ML phylogeny generation details.

**Appendix S3.** Gene Synteny, Homeolog, and Duplication analysis.

**Appendix S4.** Protein MEME Motif and structure prediction analysis.

**Appendix S5.** Promoter element mining.

**Appendix S6.** Transposable element reporting and summary.

**Appendix S7**. Extra notes Group B.

**Figure S1.** Schema outlining the search parameters for conducting a genome-wide search for *TaFMO*.

**Figure S2.** Expanded sub-phylogeny of Group A TaFMOs.

**Figure S3.** Expanded sub-phylogeny of Group B TaFMOs.

**Figure S4.** Expanded sub-phylogeny of Group C TaFMOs.

**Figure S5.** Consensus sequence (peptide) of 15 unique MEME motifs.

**Figure S6.** Group A *TaFMO* explored under various conditions and probed in literature.

**Figure S7.** Group C *TaFMO* explored under various conditions and probed in literature.

**Figure S8.** Plant *cis*-acting regulatory elements (CARE) detected in a 1.5 kb region upstream of start codons for all *TaFMOs* in Group A.

**Figure S9.** Plant *cis*-acting regulatory elements (CARE) detected in a 1.5 kb region upstream of start codons for all *TaFMO* in Group C.

**Figure S10.** Graphical display of the gene structure and transposable element analysis for *TaFMOs* in Group A.

**Figure S11.** Graphical display of the gene structure and transposable element analysis for *TaFMOs* in Group C.

**Table S1**. Local HMM search for all *TaFMO* candidates in RefSeqv2.1 and RefSeqv1.2, with gene number and splice variant details.

**Table S2**. List of all *FMO* candidates validated for *T. aestivum, T. urartu, H. vulare, O. sativa sp. Japonicus, A. thaliana, A. trichopoda*, and *G. sulphuraria*.

**Table S3**. Summary of types of *TaFMO* Group A, B, and C Promoter Elements (PlantCARE).

**Table S4**. Summary of number of TaFMO Group A, B, and C Promoter Element (PlantCARE).

**Table S5**. RNA-sequencing datasets used for Expression Profiling (Transcriptome) from WheatExpression Database (http://www.wheat-expression.com/)

**Table S6**. Literature evidence of all 170 *TaFMO*.

**Table S7**. *TaFMO* Transcriptomic (FPKM) Metadata in five tissue types obtained from the Wheat Proteome Database.

**Table S8**. TaFMO Proteomics Metadata obtained from the Wheat Proteome Database (values in spectral counts).

## Author Contributions

SS and GB conceived the project outline, and SS conducted the *in silico* analyses. SS and GB wrote and edited the manuscript. All authors read and approved the final manuscript.

## Glossary

FMO: Flavin-containing monooxygenase
‘nc’: non-canonical
FAD: flavin adenine dinucleotide
NAD(P)H: nicotinamide adenine dinucleotide phosphate
MSA: multiple sequence alignment
FPKM: Fragments Per Kilobase of transcript per Million mapped reads
Tpm: transcripts per million

