## Supporting Information for "A bird’s-eye view: exploration of the flavin-containing monooxygenase (FMO) superfamily in common wheat"

**Appendix S1** Sources for sequence data and additional information on gene differences in *TaFMO* between IWGSC RefSeqv1.2 and RefSeqv2.1, and metadata.

### *Sources for sequence and protein data*

For common wheat (*T. aestivum*, *Ta*), genome sequence and annotation files were obtained from the Ensembl Plants database (http://plants.ensembl.org/index.html; https://ftp.ensemblgenomes.ebi.ac.uk/pub/plants/release-55/gff3/triticum_aestivum/), and the original RefSeq *T. aestivum* cv. 'Chinese Spring' gene annotation files were retrieved from IWGSC, versions 1.2 and 2.1 (https://urgi.versailles.inra.fr/download/iwgsc/IWGSC_RefSeq_Annotations/). In addition to these genomic databases, the Wheat Proteome Database (https://wheatproteome.org/ (Duncan et al., 2017)) and Uniprot Database (https://www.uniprot.org/) were also utilized. RefSeq v2.1 gene IDs in wheat were consolidated with RefSeq v1.2 for appropriate wheat candidate *FMO* genes (Table S1). Differences in *TaFMO* genes between RefSeqv1.2 and RefSeqv2.1, and other metadata pertaining to *TaFMO*s, can be found in Tables S2.
The theoretical isoelectric points (pI) and molecular weights (Mw) of all TaFMO proteins were obtained through the ExPasy server (http://web.expasy.org/compute_pi/; accessed November 2022). Additionally, subcellular localization predictions were performed using Plant-mPLoc (http://www.csbio.sjtu.edu.cn/bioinf/Cell-PLoc-2/ (Chou & Shen, 2010); accessed November 2022).

For all other plant species used in our analyses as listed in Table 1, the Ensembl Plants reference genome database was used. The Uniprot database was used to corroborate putative FMO genes for a subset of plant species: *Triticum urartu* (*Tu*), *Triticum dicoccoides* (*Td*), *Aegilops tauschii* (*Aetau*), *Hodreum vulgare* (*Hv*), *Brachypodium distachyon* (*Bd*), *Oryza sativa Japonicus* (*Osj*) , *Arabidopsis thaliana* (*At*), *Amborella trichopoda* (*AmTri*), and *Galdieria sulphuraria (Gs).* For the gymnosperm species *Picea abies* (*Pa*), treegenesdb.org was used to access peptide FASTA files for the whole genome.

### *Gene search notes*

When querying the Pfam ID* for 'FMO-like' PF00743 (Interpro: IPR020946) against the InterPro database (https://www.ebi.ac.uk/interpro), 642 different domain architectures are found for this gene family. It is not clear-cut which groups are more enriched with a higher diversity of FMO fusion proteins, as certain widely cultivated food-crop species such as banana (*Musa acuminata*), grape (*Vitis vinifera*), and soybean (*Glycine max*) have fusion FMOs, but less widely-cultivated plant species such as white jute (*Corchorus capsularis*) and the wild cutgrass *Leersia perrieri* are highly enriched with various kinds of FMOs atypical from the three most widely represented (FMO-like, FMO-like x 2, pyr3).

A local HMM search yielded, in RefSeqv2.1, 171 high-confidence (HC) and 82 low confidence (LC) genes. In RefSeqv1.2, this yielded 177 HC and 67 LC genes (Table S1). LC genes found in the HMM search were found to comprised of 'non-FMO' encoding genes or poorly annotated and truncated gene fragments. Therefore, the focus was placed on HC genes only. Similarly, the search term "Flavin monooxygenase" yielded 163 mapped and 10 unmapped putative *TaFMO* in the UniProt database. The term "Flavin-containing monooxygenase" queried against the publicly available Wheat Proteome Database (Duncan et al., 2017) yielded 220 total *TaFMO* candidates (including splice variants). From here, all possible candidates obtained from the different search methods were compiled and filtered for the core FMO motifs as reported in Figures 5ab, as summarized in Table 2 and Table S2.

Of note, TraesCS5A03G1253300.1 was included in the original analysis (Table S2), but was not found in the original HMM search, but it was comprised only of the tail end of a possible FMO (no major FAD-, NAD(P)H-binding, FATGY, or FMO-identifying motif present); in a protein MSA pile-up with all other TaFMOs, TraesCS5A03G1253300.1 did not resemble any known FMO and was highly truncated, so was excluded from downstream analysis. However, in the synteny analysis conducted (see notes S3), TraesCS5A03G1253300.1 is seen to group with TraesCS4D03G0827900.1 and TraesCS4B03G0953000.1, from group A, sub-clade A2β-3. This scenario presents the complexity in conducting thorough genome-wide searches of a largely expanded gene family, where some members may be so truncated past the point of being properly recognized as being a part of the gene of interest. This gene and the circumstance is noted in Table 3 and Table S2.

For the changes which occurred between the latest wheat reference genome (cv. Chinese Spring) RefSeqv2.1 from the previous version (RefSeqv1.2), several changes between FMOs in *Ta* were accounted for in Table S2, with most changes being single amino acid residue swaps, or the inclusion of a new N-terminus string of peptides. Between the two wheat genome versions, changes were found for 14 *TaFMO*s (that is, 8.2% of all *TaFMO*s), and 7 *TaFMOs* were no longer present in RefSesv2.1, with 3 genes from RefSeq1.2 collapsed into already existing *TaFMO* genes in Refseqv2.1. For all metadata concerning *TaFMO* see Tables S2.

Several gene families closely related to the *FMO*s repeatedly emerge erroneously as potential gene hits during HMM searches, such as *squalene monooxygenase* (*SQLE*), *amino oxidoreductase* (*AOX*), *cytochrome P450* (*CYP*), and *disulfide oxidoreductase* gene families.

* As of January 2023, the Pfam database has been decommissioned, but an archive of all information from Pfam has been consolidated in the InterPro database (https://www.ebi.ac.uk/interpro/)

### *'non-canonical' classification notes:*

Truncations and/or alterations greatly affecting the integrity of the predicted structures relative to functionally characterized AtFMO orthologs helped to further assess TaFMOs with peculiar fusion architectures. Figure S1 outlines the detailed schema used for characterizing *TaFMO* genes.

Based on literature, certain known active binding sites and motifs were deemed to be crucial for a 'canonical' FMO function. These motifs are outlined in Figure 4a and Figure 4b. As such, *TaFMO* candidates that were found to be missing, truncated, or significantly altered in one or more of the four critical documented motifs (FAD-binding, FMO-identifying, NAD(P)H-binding, and 'F/LATGY'), were annotated with the letters 'nc' (non-canonical), to denote for potential loss of a 'canonically expected' FMO function. Where residues at binding sites are slightly altered, it is not possible to determine whether the FMO function is compromised and/or deviates from the assumed conventional mode of action reported by (Eswaramoorthy et al., 2006) without experimental validation.

**Appendix S2** Alignment and ML phylogeny generation details

*Generation of Figure 2:*

For the alignment of multiple FMOs for Figure 2, MAFFT (version 7.475; (Katoh et al., 2019)) E-INS-i was utilized for the total 298 FMO amino acid sequences (including protein splice variants) from *Gs* (1), *Amtri* (18), *At* (44), *Osj* (37), and *Ta* (198); Gs was used as an outgroup, seeing as red algae has only one known FMO through HMM and Uniprot searching. The E-INS-i algorithm was chosen as a best fit model for its suitability of aligning many long sequences harbouring several conserved motifs. From here, the alignment file was put through the Homo software (version 2.0; (Jermiin, 2017)) to test for homogeneity, following instructions in the manual. The lowest p-value reported for the alignment was $9.429580e^{-05}$; this was a larger p-value than the family-wise error rate (aka the Bonferroni corrected threshold) of $1.129867e^{-06}$, indicating that the assumptions of evolution in these sequences occurring under stationary, reversible, and globally homogeneous conditions were not violated. Subsequently, this alignment file was analyzed in AliStat v1.14 (Wong et al., 2014). A summary of the Ca, Cr, Cc values were recorded. Here, the alignment was masked with a threshold of Cc >= 0.5, sites included in the masked alignment. All two alignments (unmasked, Cc >=0.5) were put through IQ-TREE (version 2.2.0) accessed in the CIPRes Science Gateway server (XSEDE) through the Mesquite Zephyr package (v 3.7; www.mesquiteproject.org). Unmasked analysis in IQ-tree to construct ML phylogeny (using Model Finder Plus) = JTT+I+G4 model based on BIC yielded more lowly-supported clades than the 50% masked analysis using the same substitution model (JTT+I+G4), and the 50% masked ML phylogeny was chosen; a consensus tree generated from 1000 Standard Boot-strapping (SBS) replicates was utilized for Figure 2. Confidently predicted protein domains from SMART (previous section) that met the cut-off threshold ($E < 1e^{-5}$) were displayed using iTOL (https://itol.embl.de/) to display protein domain structural arrangements for each candidate *TaFMO*; in cases where multiple similar domains were superimposed to a significant degree, the domain with most the significant *E*-value was selected for display.

*Generation of Figure S2:*

139 total sequences were present for sub-phylogeny Group A (GSOX); Sub-clade B1 in Group B was used for outgroup for Group A, including Group A sequences for *Tu, Hv, Osj*, and *At.* Assumptions for evolution were not violated via Homo (v2) software (lowest p-value reported for the alignment was $2.004678e^{-04}$, larger than the family-wise error rate of $5.213221e^{-06}$). Alignments were both unmasked and masked (unmasked, Cc >=0.5) using AliStat v1.14. Both masked 50% and unmasked alignment for Group A were subjected to an initial 1000 Ultra-fast bootstrapping (UFB) comparison using IQ-tree in a ML phylogeny. In cases where the phylogeny could not be sufficiently resolved (≤50% bootstrap value), a red box is indicated.

In Group A, there is a main difference when comparing unmasked and 50% masked alignments in tree construction; the orphan TaFMOs (*TraesCS1D03G0536600.1-nc, TraesCS1A03G0556900.1*) that have an ortholog both *Osj* and *Tu*, but not *Hv*, are the main reason for the poor branch support value seen (marked in red square) in Group A for Figure 2 and Figure S2. In the unmasked alignment, these candidates are represented as a polytomy for subclade A2. However, in 50% masked alignments, these orphaned *TaFMO*s are seen inside the subclade A2, as potential orthologues to all other candidates in sub-clade A2. The 50% masked alignment was used for an ML phylogeny (IQ-tree), using the JTT+G4 model according to the BIC.

*Generation of Figure S3:*
119 total sequences were present for sub-phylogeny Group B (*AtFMO1*-containing clade); 1 Osj (Cow1) and 3 4ABD TaFMO from Group C were used as an outgroup for Group B. Assumptions for evolution were not violated via Homo (v2) software (lowest p-value reported for the alignment was $8.188105e^{-05}$, larger than the family-wise error rate of $7.121493e^{-06}$). Alignments were both unmasked and masked (unmasked, Cc >=0.5) using AliStat v1.14. Both masked 50% and unmasked alignment for Group B were subjected to an initial 1000 Ultra-fast bootstrapping (UFB) comparison using IQ-tree in a ML phylogeny. In cases where the phylogeny could not be sufficiently resolved (≤50% bootstrap value), a red box is indicated. In Group B, no orphan TaFMOs were detected. Both 50% masked and unmasked alignments were more similar, and this similarity is reflected on a protein motif level where more B-group unique MEME motifs were discovered over Groups A and C TaFMO MEME motifs. The 50% masked alignment was used for an ML phylogeny (IQ-tree), using the JTT+G4 model according to the BIC, for generation of Figure 5,6, and 7.

*Generation of Figure S4:*
176 total sequences were present in the sub-phylogeny for Group C; Sub-clade B1 in Group B was used for outgroup for Group C. Assumptions for evolution not violated via Homo (v2) software (lowest p-value reported for the alignment was $2.432097e^{-05}$, larger than the family-wise error rate of $3.246753e^{-06}$). Alignments were both unmasked and masked (unmasked, Cc >=0.5) using AliStat v1.14. A masked 50% alignment for Group C was subjected to an initial 1000 Ultra-fast bootstrapping (UFB) comparison using IQ-tree in a ML phylogeny. In cases where the phylogeny could not be sufficiently resolved (≤50% bootstrap value), a red box is indicated.
In Group C, an orphan TaFMO with a protein kinase conjugate (TraesCS7D03G0088000.1) and its possible unmapped paralogue (TraesCSU03G0061100.1-nc) were present in A1. The 50% masked alignment was used for an ML phylogeny (IQ-tree), using the JTT+G4 model according to the BIC.

**Appendix S3** Gene Synteny, Homeolog, and Duplication analysis
The All-vs-all scan was performed on the MCScanX Wrapper in TBtools, which generated syntenic blocks (displayed as lines connecting one gene loci to the next) for all wheat protein sequences. TaFMO homeologs (gene ‘triads’ from each sub-genome A, B, and D) were inferred by strong branch supports from standard bootstrapping analyses (>70 was considered robust) in each of the ML sub-phylogenies for Groups A, B and C (Figures S2-4); homeolog status was further corroborated with the synteny analysis from MCScanX (Table 2). Table 2 summarizes for each group A, B, and C subclade groupings for predicted homeologs. Gene duplication events for TaFMO were reported as either directly flanking the homeolog via tandem duplication (tandem), tandemly duplicated and then separated by one or a few genes inserted between the duplicates (proximal) or separated by some event leading to extensive translocation on the same or distant chromosome (dispersed). TaFMOs arising from whole-genome duplications are denoted as (WGD), that typically share orthologs with other closely related grass species. For clades in each sub-phylogeny A, B, or C that were collapsed into polytomies due to low branch support from 1000 standard bootstrapping replicates (<50), and where synteny analysis was not sufficient to determine homeolog status confidently, TaFMO candidates were grouped into one

cluster and indicated accordingly.

**Appendix S4** Protein MEME Motif and structure prediction analysis
All TaFMO, AtFMO, and OsjFMO predicted protein structures were retrieved by inputting the amino acid sequences into the Phyre2 portal (Kelley et al., 2015). Conserved motifs as outlined in Figure 4a were then annotated manually and visualized with PyMOL; a representative TaFMO from each group A, B, and C were chosen for display in Figure 4b, including several representative ‘nc’ TaFMO protein structures. To conduct a *de novo* search for putative conserved protein motifs among all TaFMOs, HvFMOs, OsjFMOs, AtFMOs, AmTriFMOs, and PpFMOs, MEME Suite (Version 5.4.1)(Bailey et al., 2015) was used; search parameters were set to a maximum of 15 motifs, with all other default settings. Variations in the residues for specific (conserved) motifs over-represented in wheat for each sub-clade were displayed in a consensus model (Figure 4a) displayed on the 3D predicted protein models (Figure 4b).

**Appendix S5** Promoter element mining
Raw files obtained from PlantCARE are provided in a zipped folder. All *TaFMO* promoter regions from RefSeqv2.1 were obtained from TBtools GUI (GXF Sequence Extract) extracting a stretch of 1.5 kbp upstream of the start codon. Promoter elements with no description of their function or elements of unknown functions/names were discarded from further use in the promoter elements analysis, as well as elements described as core promoter elements like the ‘TATA-box’ and ‘CAAT-box’ elements. Due to the high redundancy of potential elements populating each *TaFMO* promoter region, categories for all elements were assigned based on potential biological functions and displayed in Figures 6 and Figures S8 and S9, and Table S3, S4.
Meristem-specific CAREs (Laloum et al., 2013) were found in all sub-clades in Group B, most in Group A, and mostly in sub-clade C1 for Group C. The endosperm-specific CARE, GCN4 (Wu et al., 1998), was detected mostly in sub-clade A1 for Group A, only in sub-clade B2 for Group B, and rarely in Group C. The RY-element, a seed-specific CARE (ain-Ali et al., 2021), was found dispersed in Group A and Group C, but only found in sub-clade B1 for Group B. The CARE HD-Zip 1 is involved in palisade mesophyll cell differentiation (ain-Ali et al., 2021; Sasaki et al., 2019), and is present in sub-clade A1, sub-clade B2, and present for several candidates in both sub-clades in Group C. Only one gene from each of Group B and C had motif I, a root-specific CARE (ain-Ali et al., 2021), but the xylem-specific CAREs were most ubiquitously found among all *TaFMO* promoter regions. The identified tissue-specific CAREs could be validated for their roles in transcriptional regulation through experimental evidence to complement results from the different tissue-specific elements already studied.

**Appendix S6** Transposable element reporting and summary
Using the CLARITE annotations, we retrieved Transposable Elements (TEs) data (family and location information) for all *TaFMO* genes including the 2 kb regions up- and downstream of the CDS, in the wheat RefSeqv2.1 assembly. Extra information on several *TaFMO* TE insertions:
In the diversified triad in B2β-3b, all D-copies of *TaFMO* resulting from tandem duplication events show highly variable gene structures resulting from various TE insertions, and these D-

copies exhibited lower (to no) transcriptional expression against pathogens (Figure 5 – III). In contrast, the A-copy, harboring one small TE, showed higher levels of transcription than the D-copy for *Mo*-infected wheat (Figure 5 – III); Islam et al., 2016).
In Group C, several homoeologous triads exhibited variation in gene structures due to TE insertions. The 5A, 5B, and 5D triad in sub-clade C1 (C1-4) each contained various TEs resulting in diversified gene structures, where only the A-copy was more transcriptionally active in an infection time course by the fungal pathogen *Puccinia striiformis* (*Pst*)(Figure S7 – III, IV)). In sub-clade C2 (C2-1), *TraesCS7D03G0136300.1* lacked a TE insertion in the first intron region and was transcriptionally active in various conditions explored, while the A-copy homeolog (*TraesCS7A03G0147900.1*) harboring a TE was not transcriptionally active in these same conditions (Figure S11, Figure S7 – III, IV). In the highly expanded *TaFMO* cluster in C2 located on chromosomes 2 (C2-4a), three show major TE disruption while none from the highly expanded *TaFMO* in C2-4b are impacted by TE insertion (Figure S11).

**Appendix S7** Extra notes Group B
Other *TaFMOs* of interests in sub-clade B2 include a triad from B2β-3a showing high transcriptional activity in reproductive organs of wheat, except for the D-copy of the triad (*TraesCS5D03G0799800.1*) detected with higher transcriptional activity in the leaves, harbouring an extra K-oxygenase domain (Figure 2). Peptides for this triad were detected predominantly in developing reproductive wheat organs, but also in vegetative tissue like the young coleoptiles and mature flag leaves (Figure 5 – II), with high transcriptional activity detected in the stamen and spikes at anthesis. The B-copy (*TraesCS5B03G0875600.1*) is also highly upregulated at 1 hour of drought and heat stress treatment and may be linked with early drought and heat stress responses (Figure 5 – IV). For the homoeologous triad in B2β-3b, all three D-copy *TaFMOs* were truncated (‘nc’); peptides for this triad were found mostly in the developing reproductive organs (Figure 5 – II). Only A, B, and one D-nc (TraesCS5D03G0450600.1-nc) were transcriptionally reactive at the wheat grain filling stage when wheat is being infected by *Mo* (Figure 5 – III). *TraesCS5A03G0486900.1* was also expressed in wheat seedling infected by *Zt* by 14 dpi and wheat seedling infected by *Pst* at 11 dpi, but not in uninfected plants at 14 days indicating that in addition to grain development, *TraesCS5A03G0486900.1* could be responding to pathogenesis on wheat at later timepoints. A *TaFMO* in sub-clade B2β-4c(2), *TraesCS7A03G1341700.1*, is involved in total tiller number increase (Yu et al., 2021).

**Supporting Figures**

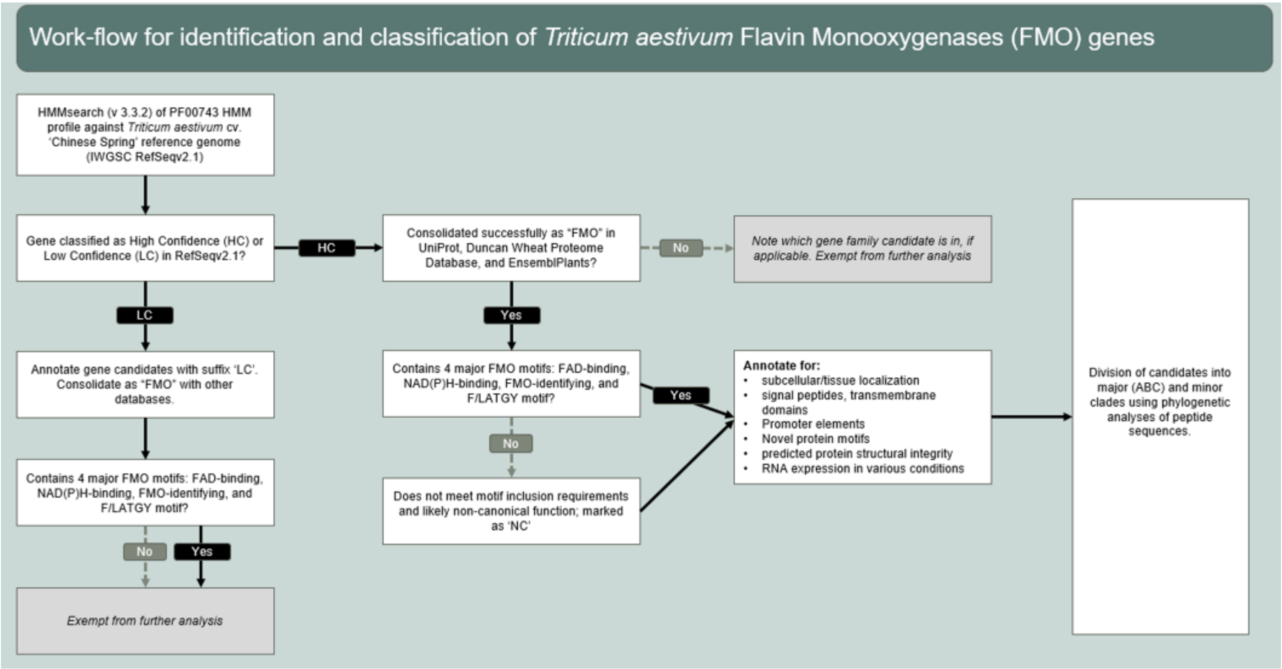


**Figure S1** Schema outlining the search parameters for conducting a genome-wide search for *TaFMO*.

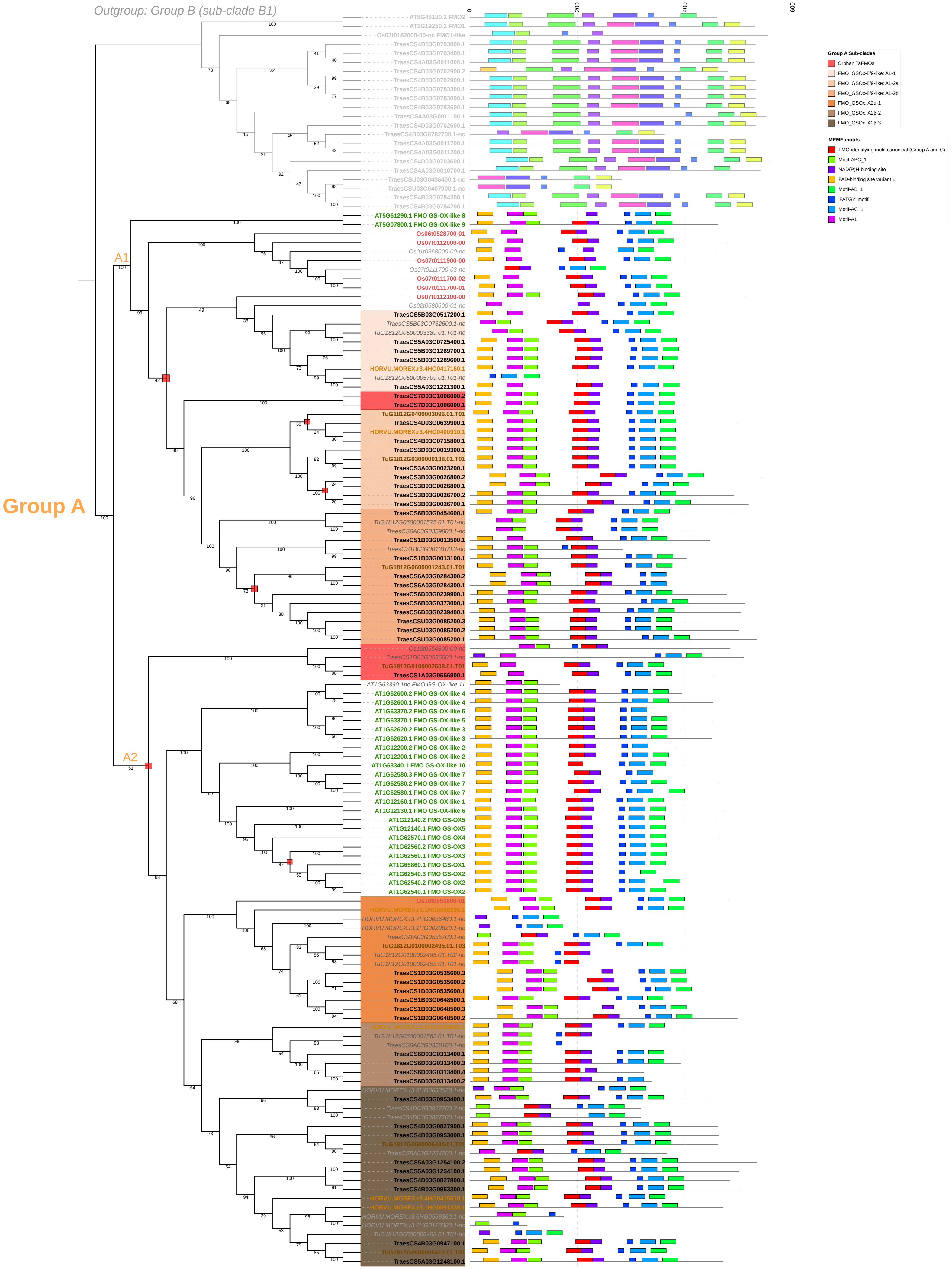
Outgroup: Group B (sub-clade B1)
Group A
A1
A2
0
200
400
600
AT5G45180.1 FMO2
AT1G19250.1 FMO1
Os03t0182000-00-nc FMO1-like
TraesCS4D03G0703000.1
TraesCS4D03G0703400.1
TraesCS4A03G0011000.1
TraesCS4D03G0702900.2
TraesCS4D03G0702900.1
TraesCS4B03G0783300.1
TraesCS4B03G0783000.1
TraesCS4B03G0783600.1
TraesCS4A03G0011100.1
TraesCS4D03G0702600.1
TraesCS4B03G0782700.1-nc
TraesCS4A03G0011700.1
TraesCS4A03G0011200.1
TraesCS4D03G0703600.1
TraesCS4A03G0010700.1
TraesCSU03G0436400.1-nc
TraesCSU03G0407900.1-nc
TraesCS4B03G0784300.1
TraesCS4B03G0784200.1
AT5G61290.1 FMO GS-OX-like 8
AT5G07800.1 FMO GS-OX-like 9
Os06t0528700-01
Os07t0112000-00
Os01t0368000-00-nc
Os07t0111900-00
Os07t0111700-03-nc
Os07t0111700-02
Os07t0111700-01
Os07t0112100-00
Os02t0580600-01-nc
TraesCS5B03G0517200.1
TraesCS5B03G0762600.1-nc
TuG1812G0500003389.01.T01-nc
TraesCS5A03G0725400.1
TraesCS5B03G1289700.1
TraesCS5B03G1289600.1
HORVU.MOREX.r3.4HG0417160.1
TuG1812G0500005709.01.T01-nc
TraesCS5A03G1221300.1
TraesCS7D03G1006000.2
TraesCS7D03G1006000.1
TuG1812G0400003096.01.T01
TraesCS4D03G0639900.1
HORVU.MOREX.r3.4HG0400910.1
TraesCS4B03G0715800.1
TraesCS3D03G0019300.1
TuG1812G0300000138.01.T01
TraesCS3A03G0023200.1
TraesCS3B03G0026800.2
TraesCS3B03G0026800.1
TraesCS3B03G0026700.2
TraesCS3B03G0026700.1
TraesCS6B03G0454600.1
TuG1812G0600001575.01.T01-nc
TraesCS6A03G0359800.1-nc
TraesCS1B03G0013500.1
TraesCS1B03G0013100.2-nc
TraesCS1B03G0013100.1
TuG1812G0600001243.01.T01
TraesCS6A03G0284300.2
TraesCS6A03G0284300.1
TraesCS6D03G0239900.1
TraesCS6B03G0373000.1
TraesCS6D03G0239400.1
TraesCSU03G0085200.3
TraesCSU03G0085200.2
TraesCSU03G0085200.1
Os10t0554300-00-nc
TraesCS1D03G0536600.1-nc
TuG1812G0100002508.01.T01
TraesCS1A03G0556900.1
AT1G63390.1nc FMO GS-OX-like 11
AT1G62600.2 FMO GS-OX-like 4
AT1G62600.1 FMO GS-OX-like 4
AT1G63370.2 FMO GS-OX-like 5
AT1G63370.1 FMO GS-OX-like 5
AT1G62620.2 FMO GS-OX-like 3
AT1G62620.1 FMO GS-OX-like 3
AT1G12200.2 FMO GS-OX-like 2
AT1G12200.1 FMO GS-OX-like 2
AT1G63340.1 FMO GS-OX-like 10
AT1G62580.3 FMO GS-OX-like 7
AT1G62580.2 FMO GS-OX-like 7
AT1G62580.1 FMO GS-OX-like 7
AT1G12160.1 FMO GS-OX-like 1
AT1G12130.1 FMO GS-OX-like 6
AT1G12140.2 FMO GS-OX5
AT1G12140.1 FMO GS-OX5
AT1G62570.1 FMO GS-OX4
AT1G62560.2 FMO GS-OX3
AT1G62560.1 FMO GS-OX3
AT1G65860.1 FMO GS-OX1
AT1G62540.3 FMO GS-OX2
AT1G62540.2 FMO GS-OX2
AT1G62540.1 FMO GS-OX2
Os10t0553800-01
HORVU.MOREX.r3.1HG0056290.1
HORVU.MOREX.r3.7HG0656460.1-nc
HORVU.MOREX.r3.1HG0029820.1-nc
TraesCS1A03G0555700.1-nc
TuG1812G0100002495.01.T03
TuG1812G0100002495.01.T02-nc
TuG1812G0100002495.01.T01-nc
TraesCS1D03G0535600.3
TraesCS1D03G0535600.2
TraesCS1D03G0535600.1
TraesCS1B03G0648500.1
TraesCS1B03G0648500.3
TraesCS1B03G0648500.2
HORVU.MOREX.r3.6HG0569600.1
TuG1812G0600001563.01.T01-nc
TraesCS6A03G0358100.1-nc
TraesCS6D03G0313400.1
TraesCS6D03G0313400.3
TraesCS6D03G0313400.4
TraesCS6D03G0313400.2
HORVU.MOREX.r3.6HG0633520.1-nc
TraesCS4B03G0953400.1
TraesCS4D03G0827700.2-nc
TraesCS4D03G0827700.1-nc
TraesCS4D03G0827900.1
TraesCS4B03G0953000.1
TuG1812G0500005494.01.T01
TraesCS5A03G1254200.1-nc
TraesCS5A03G1254100.2
TraesCS5A03G1254100.1
TraesCS4D03G0827800.1
TraesCS4B03G0953300.1
HORVU.MOREX.r3.4HG0415610.1
HORVU.MOREX.r3.1HG0091330.1
HORVU.MOREX.r3.6HG0599360.1-nc
HORVU.MOREX.r3.2HG0120380.1-nc
TuG1812G0500005493.01.T01-nc
TraesCS4B03G0947100.1
TuG1812G0500005412.01.T01
TraesCS5A03G1248100.1
Group A Sub-clades
Orphan TaFMOs
FMO_GSOx-8/9-like: A1-1
FMO_GSOx-8/9-like: A1-2a
FMO_GSOx-8/9-like: A1-2b
FMO_GSOx: A2α-1
FMO_GSOx: A2β-2
FMO_GSOx: A2β-3
MEME motifs
FMO-identifying motif canonical (Group A and C)
Motif-ABC_1
NAD(P)H-binding site
FAD-binding site variant 1
Motif-AB_1
'FATGY' motif
Motif-AC_1
Motif-A1

**Figure S2** Expanded sub-phylogeny of Group A *TaFMO*s. Maximum-likelihood consensus phylogeny (IQ-TREE, 1000 standard bootstrap replicates, JTT+G4) of all Group A TaFMO, TuFMO, HvFMO, OsjFMO, and AtFMO protein sequences. 'NC' FMOs of all species annotated with italicised and grey text. Nodes with low bootstrap support representing polytomies are marked with a red box. Confidently supported subclades for *TaFMO* are coloured accordingly in shades orange to brown, and subclade names and colours can be found in the legend and Table 2. MEME motifs with corresponding locations of motifs are displayed to the right of each FMO candidate indicating 15 of the most conserved protein motifs; scale bars indicate the length of the protein sequence in amino acids numbers. Outgroup indicated as a subclade from group B.

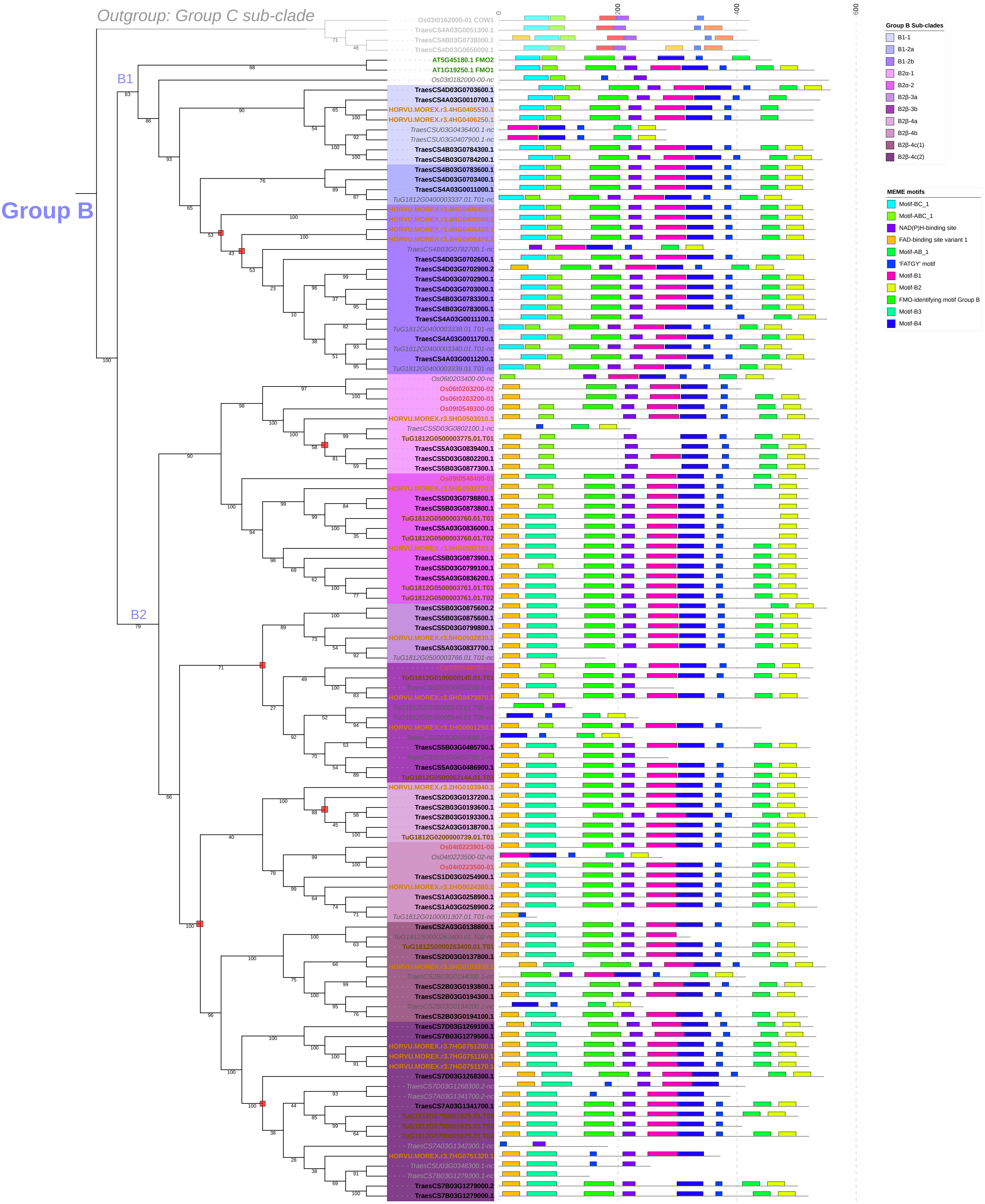
Outgroup: Group C sub-clade
Group B
B1
B2
0
200
400
600
Os03t0162000-01 COW1
TraesCS4A03G0051300.1
TraesCS4B03G0738000.1
TraesCS4D03G0656000.1
AT5G45180.1 FMO2
AT1G19250.1 FMO1
Os03t0182000-00-nc
TraesCS4D03G0703600.1
TraesCS4A03G0010700.1
HORVU.MOREX.r3.4HG0405530.1
HORVU.MOREX.r3.4HG0406250.1
TraesCSU03G0436400.1-nc
TraesCSU03G0407900.1-nc
TraesCS4B03G0784300.1
TraesCS4B03G0784200.1
TraesCS4B03G0783600.1
TraesCS4D03G0703400.1
TraesCS4A03G0011000.1
TuG1812G0400003337.01.T01-nc
HORVU.MOREX.r3.4HG0405450.1
HORVU.MOREX.r3.4HG0405500.1
HORVU.MOREX.r3.4HG0405420.1
HORVU.MOREX.r3.4HG0405470.1
TraesCS4B03G0782700.1-nc
TraesCS4D03G0702600.1
TraesCS4D03G0702900.2
TraesCS4D03G0702900.1
TraesCS4D03G0703000.1
TraesCS4B03G0783300.1
TraesCS4B03G0783000.1
TraesCS4A03G0011100.1
TuG1812G0400003338.01.T01-nc
TraesCS4A03G0011700.1
TuG1812G0400003340.01.T01-nc
TraesCS4A03G0011200.1
TuG1812G0400003339.01.T01-nc
Os06t0203400-00-nc
Os06t0203200-02
Os06t0203200-01
Os09t0549300-00
HORVU.MOREX.r3.5HG0503010.1
TraesCS5D03G0802100.1-nc
TuG1812G0500003775.01.T01
TraesCS5A03G0839400.1
TraesCS5D03G0802200.1
TraesCS5B03G0877300.1
Os09t0548400-01
HORVU.MOREX.r3.5HG0502770.1
TraesCS5D03G0798800.1
TraesCS5B03G0873800.1
TuG1812G0500003760.01.T01
TraesCS5A03G0836000.1
TuG1812G0500003760.01.T02
HORVU.MOREX.r3.5HG0502760.1
TraesCS5B03G0873900.1
TraesCS5D03G0799100.1
TraesCS5A03G0836200.1
TuG1812G0500003761.01.T01
TuG1812G0500003761.01.T02
TraesCS5B03G0875600.2
TraesCS5B03G0875600.1
TraesCS5D03G0799800.1
HORVU.MOREX.r3.5HG0502830.1
TraesCS5A03G0837700.1
TuG1812G0500003766.01.T01-nc
Os09t0548700-00
TuG1812G0100000145.01.T01
TraesCS5D03G0450200.1-nc
HORVU.MOREX.r3.5HG0473870.1
TuG1812G0100000143.01.T01-nc
TuG1812G0100000144.01.T01-nc
HORVU.MOREX.r3.1HG0001250.1
TraesCS5D03G0450600.1-nc
TraesCS5B03G0485700.1
TraesCS5D03G0450700.1-nc
TraesCS5A03G0486900.1
TuG1812G0500002144.01.T01
HORVU.MOREX.r3.2HG0103940.1
TraesCS2D03G0137200.1
TraesCS2B03G0193600.1
TraesCS2B03G0193300.1
TraesCS2A03G0138700.1
TuG1812G0200000739.01.T01
Os04t0223901-00
Os04t0223500-02-nc
Os04t0223500-01
TraesCS1D03G0254900.1
HORVU.MOREX.r3.1HG0024380.1
TraesCS1A03G0258900.1
TraesCS1A03G0258900.2
TuG1812G0100001307.01.T01-nc
TraesCS2A03G0138800.1
TuG1812S0000263400.01.T02-nc
TuG1812S0000263400.01.T01
TraesCS2D03G0137800.1
HORVU.MOREX.r3.2HG0103930.1
TraesCS2B03G0194000.1-nc
TraesCS2B03G0193800.1
TraesCS2B03G0194300.1
TraesCS2B03G0194200.1-nc
TraesCS2B03G0194100.1
TraesCS7D03G1269100.1
TraesCS7B03G1279500.1
HORVU.MOREX.r3.7HG0751200.1
HORVU.MOREX.r3.7HG0751160.1
HORVU.MOREX.r3.7HG0751170.1
TraesCS7D03G1268300.1
TraesCS7D03G1268300.2-nc
TraesCS7A03G1341700.2-nc
TraesCS7A03G1341700.1
TuG1812G0700005925.01.T03
TuG1812G0700005925.01.T01
TuG1812G0700005925.01.T02
TraesCS7A03G1342300.1-nc
HORVU.MOREX.r3.7HG0751320.1
TraesCSU03G0348300.1-nc
TraesCS7B03G1279300.1-nc
TraesCS7B03G1279000.2
TraesCS7B03G1279000.1
Group B Sub-clades
B1-1
B1-2a
B1-2b
B2α-1
B2α-2
B2β-3a
B2β-3b
B2β-4a
B2β-4b
B2β-4c(1)
B2β-4c(2)
MEME motifs
Motif-BC_1
Motif-ABC_1
NAD(P)H-binding site
FAD-binding site variant 1
Motif-AB_1
'FATGY' motif
Motif-B1
Motif-B2
FMO-identifying motif Group B
Motif-B3
Motif-B4

**Figure S3** Expanded sub-phylogeny of Group B *TaFMO*s. Maximum-likelihood consensus phylogeny (IQ-TREE, 1000 standard bootstrap replicates, JTT+G4) of all Group B TaFMO, TuFMO, HvFMO, OsjFMO, and AtFMO protein sequences. ‘NC’ FMOs of all species annotated with italicised and grey text. Nodes with low bootstrap support representing polytomies are marked with a red box. Confidently supported subclades for TaFMO are coloured accordingly in shades of purple to pink, and subclade names and colours can be found in the legend and Table 2. MEME motifs with corresponding locations of motifs are displayed to the right of each FMO candidate indicating 15 of the most conserved protein motifs; scale bars indicate the length of the protein sequence in amino acids numbers. Outgroup indicated as a subclade from group C.

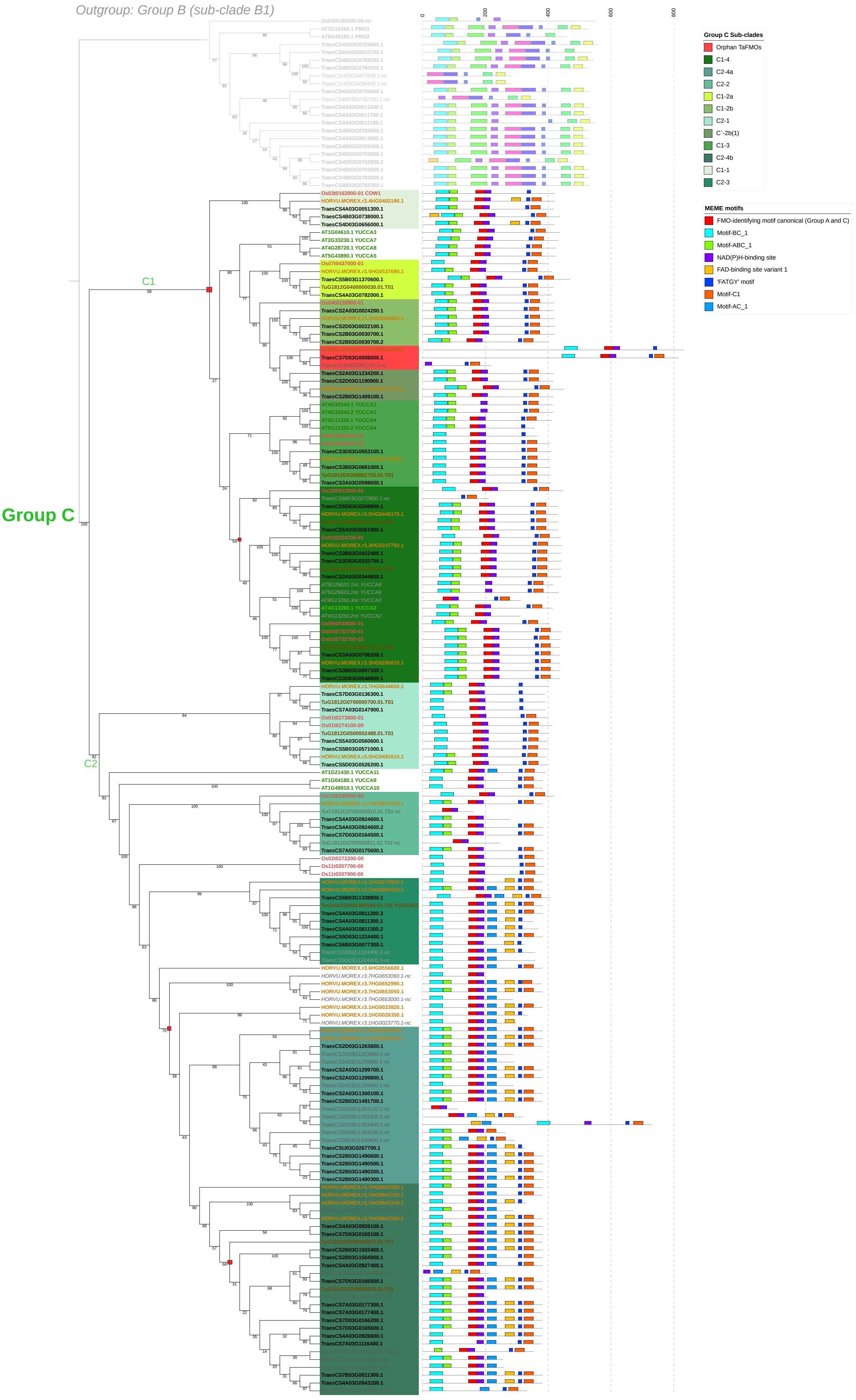
Outgroup: Group B (sub-clade B1)
Group C
C1
C2
Group C Sub-clades
Orphan TaFMOs
C1-4
C2-4a
C2-2
C1-2a
C1-2b
C2-1
C`-2b(1)
C1-3
C2-4b
C1-1
C2-3
MEME motifs
FMO-identifying motif canonical (Group A and C)
Motif-BC_1
Motif-ABC_1
NAD(P)H-binding site
FAD-binding site variant 1
'FATGY' motif
Motif-C1
Motif-AC_1

**Figure S4** Expanded sub-phylogeny of Group C *TaFMO*s. Maximum-likelihood consensus phylogeny (IQ-TREE, 1000 standard bootstrap replicates, JTT+G4) of all Group C TaFMO, TuFMO, HvFMO, OsjFMO, and AtFMO protein sequences. 'NC' FMOs of all species annotated with italicised and grey text. Nodes with low bootstrap support representing polytomies are marked with a red box. Confidently supported subclades for TaFMO are coloured accordingly in shades of green, and subclade names and colours can be found in the legend and Table 2. MEME motifs with corresponding locations of motifs are displayed to the right of each FMO candidate indicating 15 of the most conserved protein motifs; scale bars indicate the length of the protein sequence in amino acids numbers. Outgroup indicated as a subclade from group B

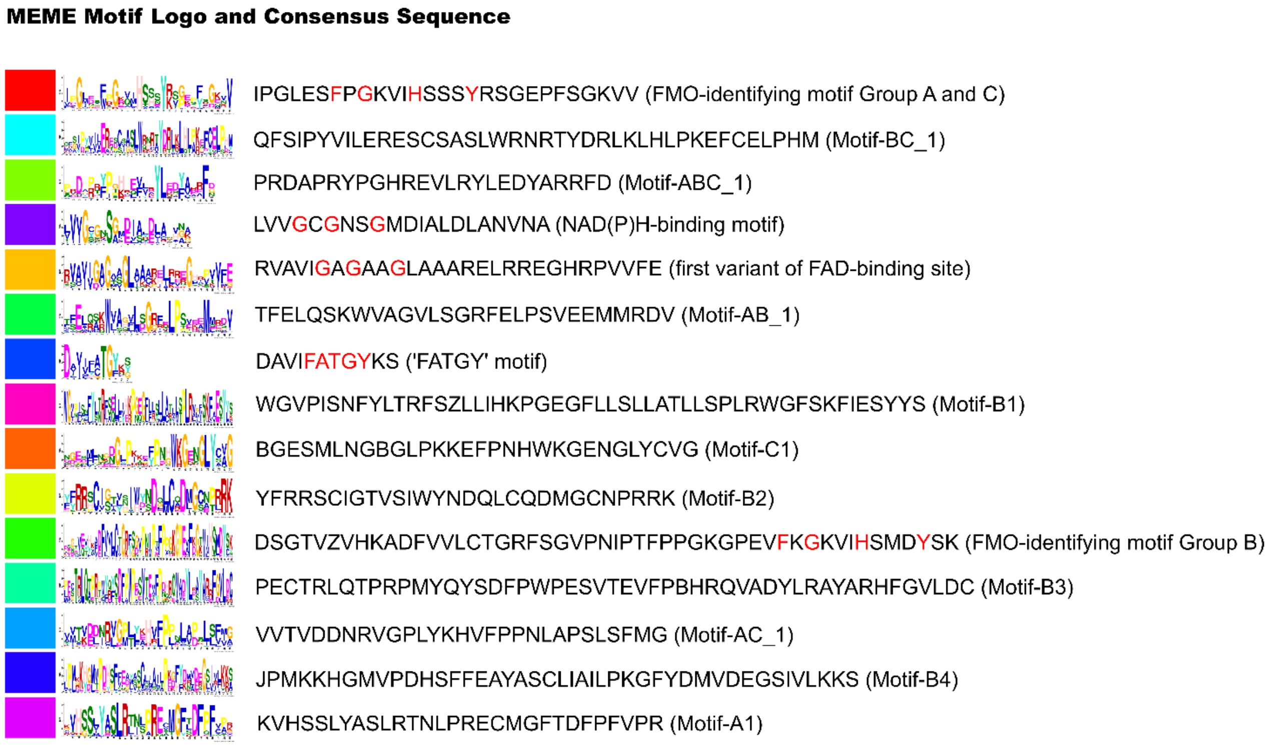


**Figure S5** Consensus sequence (peptide) of 15 unique MEME motifs. Conserved residues reported in previous literature are coloured in red.

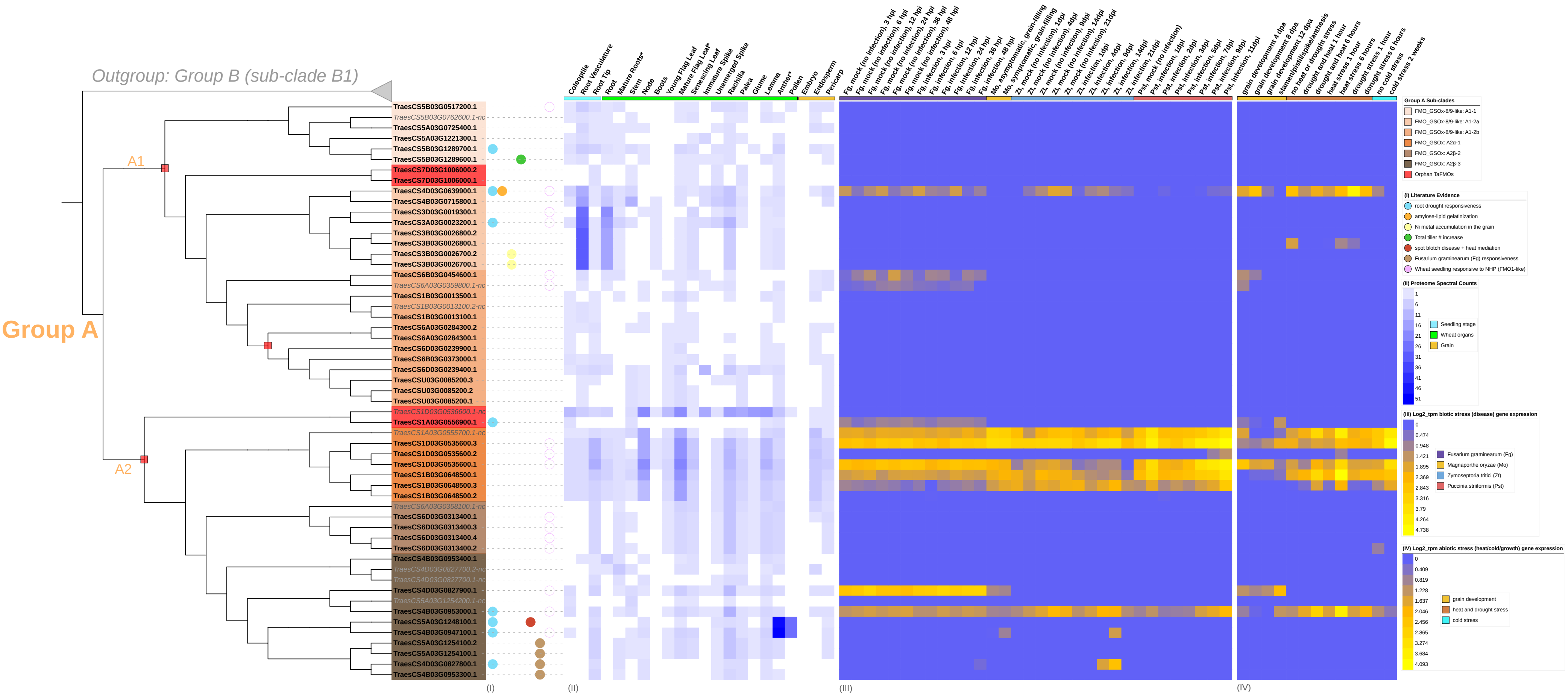

Outgroup: Group B (sub-clade B1)
Group A
A1
A2
TraesCS5B03G0517200.1
TraesCS5B03G0762600.1-nc
TraesCS5A03G0725400.1
TraesCS5A03G1221300.1
TraesCS5B03G1289700.1
TraesCS5B03G1289600.1
TraesCS7D03G1006000.2
TraesCS7D03G1006000.1
TraesCS4D03G0639900.1
TraesCS4B03G0715800.1
TraesCS3D03G0019300.1
TraesCS3A03G0023200.1
TraesCS3B03G0026800.2
TraesCS3B03G0026800.1
TraesCS3B03G0026700.2
TraesCS3B03G0026700.1
TraesCS6B03G0454600.1
TraesCS6A03G0359800.1-nc
TraesCS1B03G0013500.1
TraesCS1B03G0013100.2-nc
TraesCS1B03G0013100.1
TraesCS6A03G0284300.2
TraesCS6A03G0284300.1
TraesCS6D03G0239900.1
TraesCS6B03G0373000.1
TraesCS6D03G0239400.1
TraesCSU03G0085200.3
TraesCSU03G0085200.2
TraesCSU03G0085200.1
TraesCS1D03G0536600.1-nc
TraesCS1A03G0556900.1
TraesCS1A03G0555700.1-nc
TraesCS1D03G0535600.3
TraesCS1D03G0535600.2
TraesCS1D03G0535600.1
TraesCS1B03G0648500.1
TraesCS1B03G0648500.3
TraesCS1B03G0648500.2
TraesCS6A03G0358100.1-nc
TraesCS6D03G0313400.1
TraesCS6D03G0313400.3
TraesCS6D03G0313400.4
TraesCS6D03G0313400.2
TraesCS4B03G0953400.1
TraesCS4D03G0827700.2-nc
TraesCS4D03G0827700.1-nc
TraesCS4D03G0827900.1
TraesCS5A03G1254200.1-nc
TraesCS4B03G0953000.1
TraesCS5A03G1248100.1
TraesCS4B03G0947100.1
TraesCS5A03G1254100.2
TraesCS5A03G1254100.1
TraesCS4D03G0827800.1
TraesCS4B03G0953300.1
Coleoptile
Root Vasculature
Root Tip
Root
Mature Roots*
Stem
Node
Boots
Young Flag Leaf
Mature Flag Leaf*
Senescing Leaf
Immature Spike
Unemerged Spike
Rachilla
Palea
Glume
Lemma
Anther*
Pollen
Embryo
Endosperm
Pericarp
Fg, mock (no infection), 3 hpi
Fg, mock (no infection), 6 hpi
Fg, mock (no infection), 12 hpi
Fg, mock (no infection), 24 hpi
Fg, mock (no infection), 36 hpi
Fg, mock (no infection), 48 hpi
Fg, infection, 3 hpi
Fg, infection, 6 hpi
Fg, infection, 12 hpi
Fg, infection, 24 hpi
Fg, infection, 36 hpi
Fg, infection, 48 hpi
Mo, asymptomatic, grain-filling
Mo, symptomatic, grain-filling
Zt, mock (no infection), 1dpi
Zt, mock (no infection), 4dpi
Zt, mock (no infection), 9dpi
Zt, mock (no infection), 14dpi
Zt, mock (no infection), 21dpi
Zt, infection, 1dpi
Zt, infection, 4dpi
Zt, infection, 9dpi
Zt, infection, 14dpi
Zt, infection, 21dpi
Pst, mock (no infection)
Pst, infection, 1dpi
Pst, infection, 2dpi
Pst, infection, 3dpi
Pst, infection, 5dpi
Pst, infection, 7dpi
Pst, infection, 9dpi
Pst, infection, 11dpi
grain development 4 dpa
grain development 8 dpa
grain development 12 dpa
stamen/pistil/spike/anthesis
no heat or drought stress
drought and heat 1 hour
drought and heat 6 hours
heat stress 1 hour
heat stress 6 hours
drought stress 1 hour
drought stress 6 hours
no cold stress
cold stress 2 weeks
(I)
(II)
(III)
(IV)
Group A Sub-clades
FMO_GSOx-8/9-like: A1-1
FMO_GSOx-8/9-like: A1-2a
FMO_GSOx-8/9-like: A1-2b
FMO_GSOx: A2α-1
FMO_GSOx: A2β-2
FMO_GSOx: A2β-3
Orphan TaFMOs
(I) Literature Evidence
root drought responsiveness
amylose-lipid gelatinization
Ni metal accumulation in the grain
Total tiller # increase
spot blotch disease + heat mediation
Fusarium graminearum (Fg) responsiveness
Wheat seedling responsive to NHP (FMO1-like)
(II) Proteome Spectral Counts
1
6
11
16
21
26
31
36
41
46
51
Seedling stage
Wheat organs
Grain
(III) Log2_tpm biotic stress (disease) gene expression
0
0.474
0.948
1.421
1.895
2.369
2.843
3.316
3.79
4.264
4.738
Fusarium graminearum (Fg)
Magnaporthe oryzae (Mo)
Zymoseptoria tritici (Zt)
Puccinia striiformis (Pst)
(IV) Log2_tpm abiotic stress (heat/cold/growth) gene expression
0
0.409
0.819
1.228
1.637
2.046
2.456
2.865
3.274
3.684
4.093
grain development
heat and drought stress
cold stress

**Figure S6** Group A *TaFMO* explored under various conditions and probed in literature. (I) reports of previous evidence of transcript detection or cloning under various conditions described in the legend. (II) Proteomic expression (spectral peptide counts) for each candidate in various tissues from https://wheatproteome.org/ (Duncan et al. 2017). (III) Heatmap of *TaFMO* transcripts in response to four types of biotic stresses; response to *Fusarium graminearum* infection (*Fg*, both mock-inoculation and pathogen infection) time-course of 3, 6, 12, 24, 36, and 48 hours post infection (hpi); *Magnaporthe oryza, Mo,* fungal pathogen infection explored in asymptomatic (control) and symptomatic (infection) conditions at grain-filling stage; *Zymoseptoria tritici* (*Zt,* both mock-inoculation and pathogen infection) time-course 1, 4, 9, 14, 21 days post infection (dpi); *Puccinia striiformis* (*Pst*, one control uninfected and pathogen infection) time-course 1, 2, 3, 5, 7, 9, 11 dpi. (IV) Heatmap of *TaFMO* transcript response to grain development over a time-course (4, 8 12 days post anthesis), heat and drought stress (1 and 6 hours), and cold stress (control and 2-week cold stress). Heatmap data is displayed as Log2 transcripts per million (tpm), with blue being 0 and higher-fold expression approaching bright yellow (see legends).

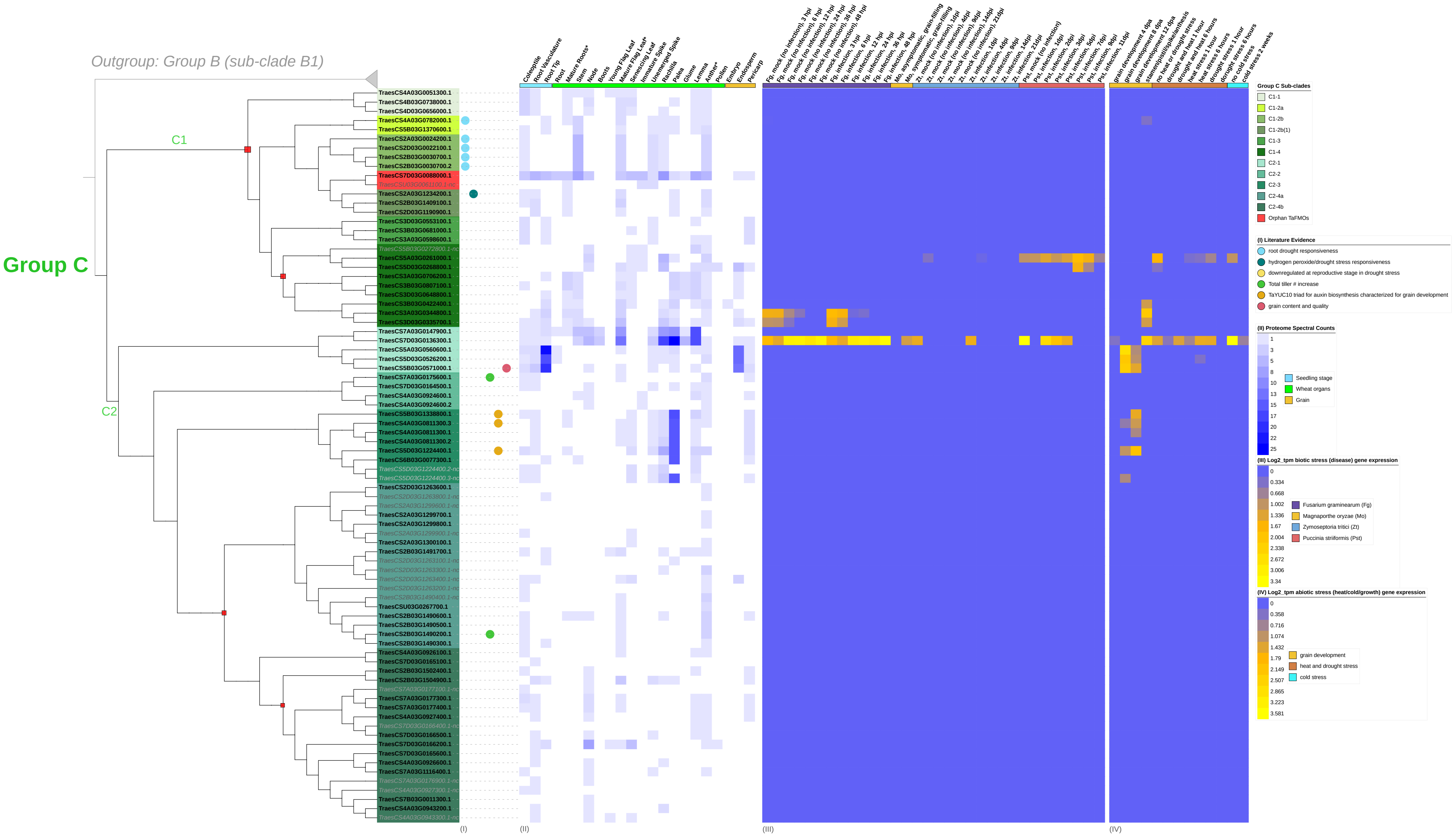

Outgroup: Group B (sub-clade B1)
Group C
C1
C2
TraesCS4A03G0051300.1
TraesCS4B03G0738000.1
TraesCS4D03G0656000.1
TraesCS4A03G0782000.1
TraesCS5B03G1370600.1
TraesCS2A03G0024200.1
TraesCS2D03G0022100.1
TraesCS2B03G0030700.1
TraesCS2B03G0030700.2
TraesCS7D03G0088000.1
TraesCSU03G0061100.1-nc
TraesCS2A03G1234200.1
TraesCS2B03G1409100.1
TraesCS2D03G1190900.1
TraesCS3D03G0553100.1
TraesCS3B03G0681000.1
TraesCS3A03G0598600.1
TraesCS5B03G0272800.1-nc
TraesCS5A03G0261000.1
TraesCS5D03G0268800.1
TraesCS3A03G0706200.1
TraesCS3B03G0807100.1
TraesCS3D03G0648800.1
TraesCS3B03G0422400.1
TraesCS3A03G0344800.1
TraesCS3D03G0335700.1
TraesCS7A03G0147900.1
TraesCS7D03G0136300.1
TraesCS5A03G0560600.1
TraesCS5D03G0526200.1
TraesCS5B03G0571000.1
TraesCS7A03G0175600.1
TraesCS7D03G0164500.1
TraesCS4A03G0924600.1
TraesCS4A03G0924600.2
TraesCS5B03G1338800.1
TraesCS4A03G0811300.3
TraesCS4A03G0811300.1
TraesCS4A03G0811300.2
TraesCS5D03G1224400.1
TraesCS6B03G0077300.1
TraesCS5D03G1224400.2-nc
TraesCS5D03G1224400.3-nc
TraesCS2D03G1263600.1
TraesCS2D03G1263800.1-nc
TraesCS2A03G1299600.1-nc
TraesCS2A03G1299700.1
TraesCS2A03G1299800.1
TraesCS2A03G1299900.1-nc
TraesCS2A03G1300100.1
TraesCS2B03G1491700.1
TraesCS2D03G1263100.1-nc
TraesCS2D03G1263300.1-nc
TraesCS2D03G1263400.1-nc
TraesCS2D03G1263200.1-nc
TraesCS2B03G1490400.1-nc
TraesCSU03G0267700.1
TraesCS2B03G1490600.1
TraesCS2B03G1490500.1
TraesCS2B03G1490200.1
TraesCS2B03G1490300.1
TraesCS4A03G0926100.1
TraesCS7D03G0165100.1
TraesCS2B03G1502400.1
TraesCS2B03G1504900.1
TraesCS7A03G0177100.1-nc
TraesCS7A03G0177300.1
TraesCS7A03G0177400.1
TraesCS4A03G0927400.1
TraesCS7D03G0166400.1-nc
TraesCS7D03G0166500.1
TraesCS7D03G0166200.1
TraesCS7D03G0165600.1
TraesCS4A03G0926600.1
TraesCS7A03G1116400.1
TraesCS7A03G0176900.1-nc
TraesCS4A03G0927300.1-nc
TraesCS7B03G0011300.1
TraesCS4A03G0943200.1
TraesCS4A03G0943300.1-nc
Coleoptile
Root Vasculature
Root Tip
Root
Mature Roots*
Stem
Node
Boots
Young Flag Leaf
Mature Flag Leaf*
Senescing Leaf
Immature Spike
Unemerged Spike
Rachilla
Palea
Glume
Lemma
Anther*
Pollen
Embryo
Endosperm
Pericarp
Fg, mock (no infection), 3 hpi
Fg, mock (no infection), 6 hpi
Fg, mock (no infection), 12 hpi
Fg, mock (no infection), 24 hpi
Fg, mock (no infection), 36 hpi
Fg, mock (no infection), 48 hpi
Fg, infection, 3 hpi
Fg, infection, 6 hpi
Fg, infection, 12 hpi
Fg, infection, 24 hpi
Fg, infection, 36 hpi
Fg, infection, 48 hpi
Mo, asymptomatic, grain-filling
Mo, symptomatic, grain-filling
Zt, mock (no infection), 1dpi
Zt, mock (no infection), 4dpi
Zt, mock (no infection), 9dpi
Zt, mock (no infection), 14dpi
Zt, mock (no infection), 21dpi
Zt, infection, 1dpi
Zt, infection, 4dpi
Zt, infection, 9dpi
Zt, infection, 14dpi
Zt, infection, 21dpi
Pst, mock (no infection)
Pst, infection, 1dpi
Pst, infection, 2dpi
Pst, infection, 3dpi
Pst, infection, 5dpi
Pst, infection, 7dpi
Pst, infection, 9dpi
Pst, infection, 11dpi
grain development 4 dpa
grain development 8 dpa
grain development 12 dpa
stamen/pistil/spike/anthesis
no heat or drought stress
drought and heat 1 hour
drought and heat 6 hours
heat stress 1 hour
heat stress 6 hours
drought stress 1 hour
dorught stress 6 hours
no cold stress
cold stress 2 weeks
(I)
(II)
(III)
(IV)
Group C Sub-clades
C1-1
C1-2a
C1-2b
C1-2b(1)
C1-3
C1-4
C2-1
C2-2
C2-3
C2-4a
C2-4b
Orphan TaFMOs
(I) Literature Evidence
root drought responsiveness
hydrogen peroxide/drought stress responsiveness
downregulated at reproductive stage in drought stress
Total tiller # increase
TaYUC10 triad for auxin biosynthesis characterized for grain development
grain content and quality
(II) Proteome Spectral Counts
1
3
5
8
10
13
15
17
20
22
25
Seedling stage
Wheat organs
Grain
(III) Log2_tpm biotic stress (disease) gene expression
0
0.334
0.668
1.002
1.336
1.67
2.004
2.338
2.672
3.006
3.34
Fusarium graminearum (Fg)
Magnaporthe oryzae (Mo)
Zymoseptoria tritici (Zt)
Puccinia striiformis (Pst)
(IV) Log2_tpm abiotic stress (heat/cold/growth) gene expression
0
0.358
0.716
1.074
1.432
1.79
2.149
2.507
2.865
3.223
3.581
grain development
heat and drought stress
cold stress

**Figure S7** Group C *TaFMO* explored under various conditions and probed in literature. (I) reports of previous evidence of transcript detection or cloning under various conditions described in the legend. (II) Proteomic expression (spectral peptide counts) for each candidate in various tissues from https://wheatproteome.org/ (Duncan et al. 2017). (III) Heatmap of *TaFMO* transcripts in response to four types of biotic stresses; response to *Fusarium graminearum* infection (*Fg*, both mock-inoculation and pathogen infection) time-course of 3, 6, 12, 24, 36, and 48 hours post infection (hpi); *Magnaporthe oryza, Mo,* fungal pathogen infection explored in asymptomatic (control) and symptomatic (infection) conditions at grain-filling stage; *Zymoseptoria tritici* (*Zt,* both mock-inoculation and pathogen infection) time-course 1, 4, 9, 14, 21 days post infection (dpi); *Puccinia striiformis* (*Pst*, one control uninfected and pathogen infection) time-course 1, 2, 3, 5, 7, 9, 11 dpi. (IV) Heatmap of *TaFMO* transcript response to grain development over a time-course (4, 8 12 days post anthesis), heat and drought stress (1 and 6 hours), and cold stress (control and 2-week cold stress). Heatmap data is displayed as Log2 transcripts per million (tpm), with blue being 0 and higher-fold expression approaching bright yellow (see legends).

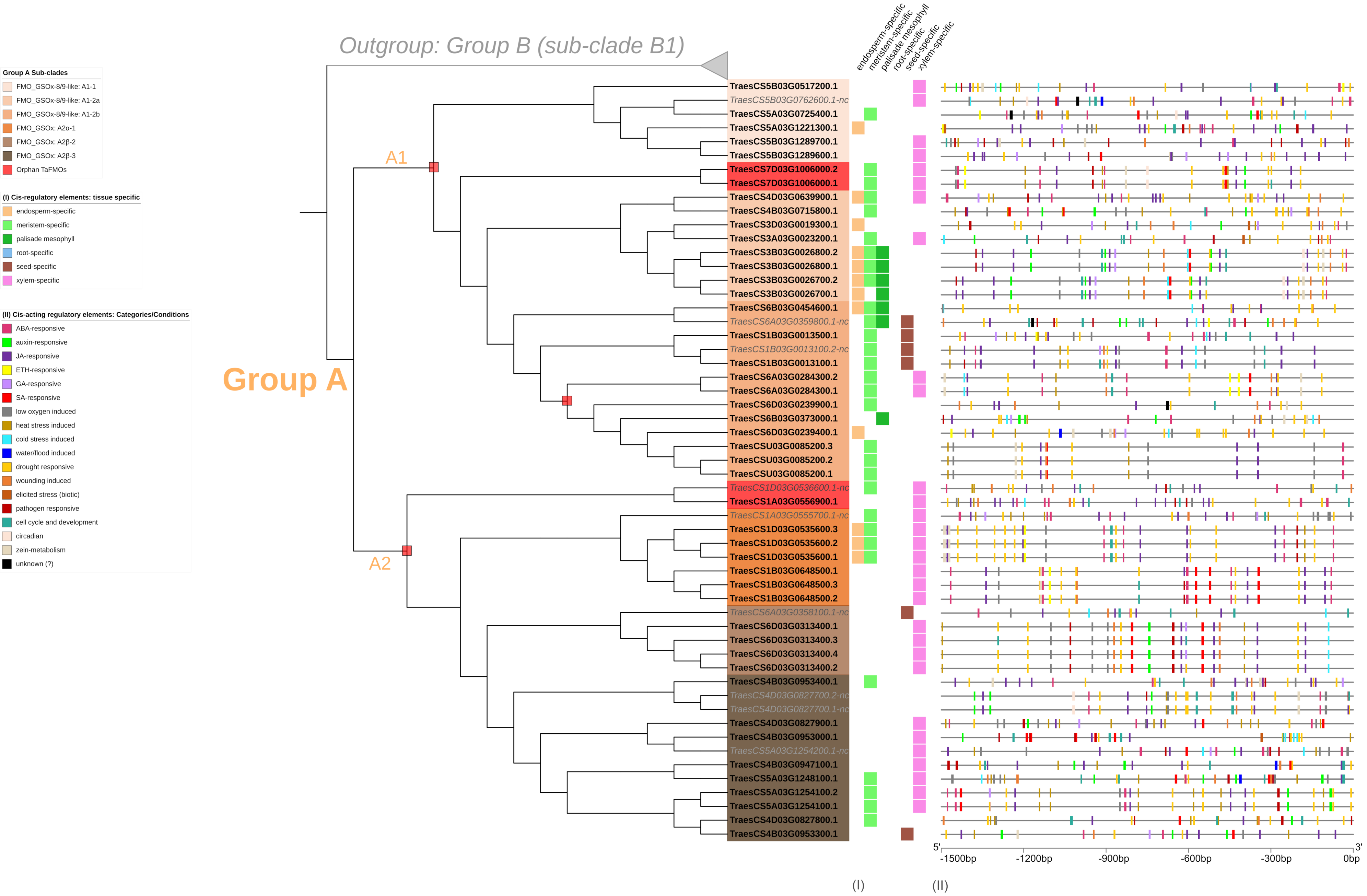

Outgroup: Group B (sub-clade B1)
Group A
A1
A2
Group A Sub-clades
FMO_GSOx-8/9-like: A1-1
FMO_GSOx-8/9-like: A1-2a
FMO_GSOx-8/9-like: A1-2b
FMO_GSOx: A2α-1
FMO_GSOx: A2β-2
FMO_GSOx: A2β-3
Orphan TaFMOs
(I) Cis-regulatory elements: tissue specific
endosperm-specific
meristem-specific
palisade mesophyll
root-specific
seed-specific
xylem-specific
(II) Cis-acting regulatory elements: Categories/Conditions
ABA-responsive
auxin-responsive
JA-responsive
ETH-responsive
GA-responsive
SA-responsive
low oxygen induced
heat stress induced
cold stress induced
water/flood induced
drought responsive
wounding induced
elicited stress (biotic)
pathogen responsive
cell cycle and development
circadian
zein-metabolism
unknown (?)
TraesCS5B03G0517200.1
TraesCS5B03G0762600.1-nc
TraesCS5A03G0725400.1
TraesCS5A03G1221300.1
TraesCS5B03G1289700.1
TraesCS5B03G1289600.1
TraesCS7D03G1006000.2
TraesCS7D03G1006000.1
TraesCS4D03G0639900.1
TraesCS4B03G0715800.1
TraesCS3D03G0019300.1
TraesCS3A03G0023200.1
TraesCS3B03G0026800.2
TraesCS3B03G0026800.1
TraesCS3B03G0026700.2
TraesCS3B03G0026700.1
TraesCS6B03G0454600.1
TraesCS6A03G0359800.1-nc
TraesCS1B03G0013500.1
TraesCS1B03G0013100.2-nc
TraesCS1B03G0013100.1
TraesCS6A03G0284300.2
TraesCS6A03G0284300.1
TraesCS6D03G0239900.1
TraesCS6B03G0373000.1
TraesCS6D03G0239400.1
TraesCSU03G0085200.3
TraesCSU03G0085200.2
TraesCSU03G0085200.1
TraesCS1D03G0536600.1-nc
TraesCS1A03G0556900.1
TraesCS1A03G0555700.1-nc
TraesCS1D03G0535600.3
TraesCS1D03G0535600.2
TraesCS1D03G0535600.1
TraesCS1B03G0648500.1
TraesCS1B03G0648500.3
TraesCS1B03G0648500.2
TraesCS6A03G0358100.1-nc
TraesCS6D03G0313400.1
TraesCS6D03G0313400.3
TraesCS6D03G0313400.4
TraesCS6D03G0313400.2
TraesCS4B03G0953400.1
TraesCS4D03G0827700.2-nc
TraesCS4D03G0827700.1-nc
TraesCS4D03G0827900.1
TraesCS4B03G0953000.1
TraesCS5A03G1254200.1-nc
TraesCS4B03G0947100.1
TraesCS5A03G1248100.1
TraesCS5A03G1254100.2
TraesCS5A03G1254100.1
TraesCS4D03G0827800.1
TraesCS4B03G0953300.1
5'
3'
-1500bp
-1200bp
-900bp
-600bp
-300bp
0bp
(I)
(II)

**Figure S8** Plant *cis*-acting regulatory elements (CARE) detected in a 1.5 kb region upstream of start codons for all *TaFMOs* in Group A. X-axis denote the number of base pairs away from the start codon, where 5’ is 1,500 bp upstream ATG start codon (3’ and 0 bp is right before the ATG start codon). Polytomy regions are denoted by a red box. The presence of regulatory elements specific to different tissues are described in (I). Other responsive promoter elements are denoted in the figure legend (II), and displayed as rectangles of corresponding color to the condition in (II).

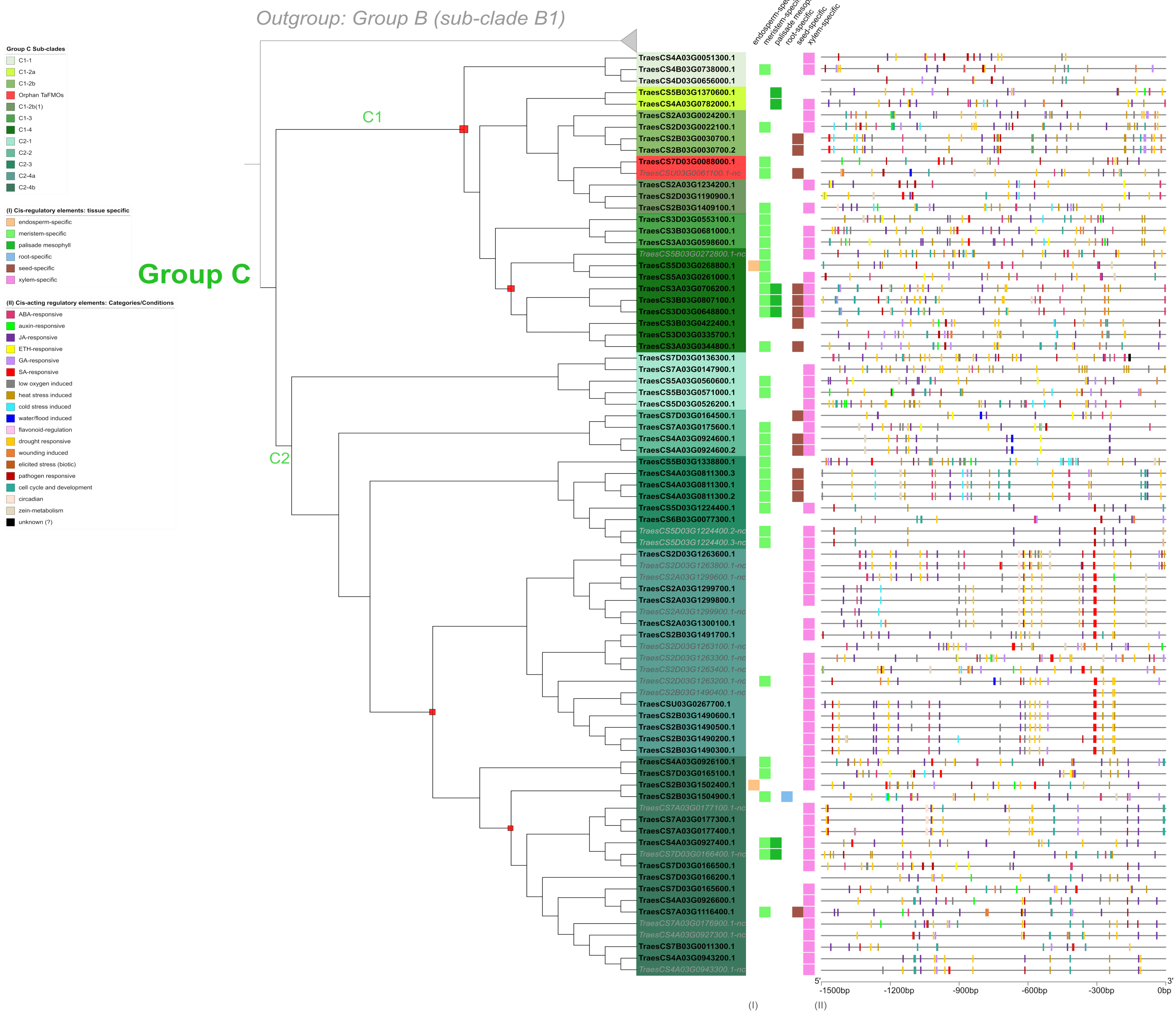

Outgroup: Group B (sub-clade B1)
Group C Sub-clades
C1-1
C1-2a
C1-2b
Orphan TaFMOs
C1-2b(1)
C1-3
C1-4
C2-1
C2-2
C2-3
C2-4a
C2-4b
(I) Cis-regulatory elements: tissue specific
endosperm-specific
meristem-specific
palisade mesophyll
root-specific
seed-specific
xylem-specific
(II) Cis-acting regulatory elements: Categories/Conditions
ABA-responsive
auxin-responsive
JA-responsive
ETH-responsive
GA-responsive
SA-responsive
low oxygen induced
heat stress induced
cold stress induced
water/flood induced
flavonoid-regulation
drought responsive
wounding induced
elicited stress (biotic)
pathogen responsive
cell cycle and development
circadian
zein-metabolism
unknown (?)
Group C
C1
C2
TraesCS4A03G0051300.1
TraesCS4B03G0738000.1
TraesCS4D03G0656000.1
TraesCS5B03G1370600.1
TraesCS4A03G0782000.1
TraesCS2A03G0024200.1
TraesCS2D03G0022100.1
TraesCS2B03G0030700.1
TraesCS2B03G0030700.2
TraesCS7D03G0088000.1
TraesCSU03G0061100.1-nc
TraesCS2A03G1234200.1
TraesCS2D03G1190900.1
TraesCS2B03G1409100.1
TraesCS3D03G0553100.1
TraesCS3B03G0681000.1
TraesCS3A03G0598600.1
TraesCS5B03G0272800.1-nc
TraesCS5D03G0268800.1
TraesCS5A03G0261000.1
TraesCS3A03G0706200.1
TraesCS3B03G0807100.1
TraesCS3D03G0648800.1
TraesCS3B03G0422400.1
TraesCS3D03G0335700.1
TraesCS3A03G0344800.1
TraesCS7D03G0136300.1
TraesCS7A03G0147900.1
TraesCS5A03G0560600.1
TraesCS5B03G0571000.1
TraesCS5D03G0526200.1
TraesCS7D03G0164500.1
TraesCS7A03G0175600.1
TraesCS4A03G0924600.1
TraesCS4A03G0924600.2
TraesCS5B03G1338800.1
TraesCS4A03G0811300.3
TraesCS4A03G0811300.1
TraesCS4A03G0811300.2
TraesCS5D03G1224400.1
TraesCS6B03G0077300.1
TraesCS5D03G1224400.2-nc
TraesCS5D03G1224400.3-nc
TraesCS2D03G1263600.1
TraesCS2D03G1263800.1-nc
TraesCS2A03G1299600.1-nc
TraesCS2A03G1299700.1
TraesCS2A03G1299800.1
TraesCS2A03G1299900.1-nc
TraesCS2A03G1300100.1
TraesCS2B03G1491700.1
TraesCS2D03G1263100.1-nc
TraesCS2D03G1263300.1-nc
TraesCS2D03G1263400.1-nc
TraesCS2D03G1263200.1-nc
TraesCS2B03G1490400.1-nc
TraesCSU03G0267700.1
TraesCS2B03G1490600.1
TraesCS2B03G1490500.1
TraesCS2B03G1490200.1
TraesCS2B03G1490300.1
TraesCS4A03G0926100.1
TraesCS7D03G0165100.1
TraesCS2B03G1502400.1
TraesCS2B03G1504900.1
TraesCS7A03G0177100.1-nc
TraesCS7A03G0177300.1
TraesCS7A03G0177400.1
TraesCS4A03G0927400.1
TraesCS7D03G0166400.1-nc
TraesCS7D03G0166500.1
TraesCS7D03G0166200.1
TraesCS7D03G0165600.1
TraesCS4A03G0926600.1
TraesCS7A03G1116400.1
TraesCS7A03G0176900.1-nc
TraesCS4A03G0927300.1-nc
TraesCS7B03G0011300.1
TraesCS4A03G0943200.1
TraesCS4A03G0943300.1-nc
endosperm-specific
meristem-specific
palisade mesophyll
root-specific
seed-specific
xylem-specific
5'
3'
-1500bp
-1200bp
-900bp
-600bp
-300bp
0bp
(I)
(II)

**Figure S9** Plant *cis*-acting regulatory elements (CARE) detected in a 1.5 kb region upstream of start codons for all *TaFMOs* in Group C. X-axis denote the number of base pairs away from the start codon, where 5' is 1,500 bp upstream ATG start codon (3' and 0 bp is right before the ATG start codon). Polytomy regions are denoted by a red box. The presence of regulatory elements specific to different tissues are described in (I). Other responsive promoter elements are denoted in the figure legend (II), and displayed as rectangles of corresponding color to the condition in (II).

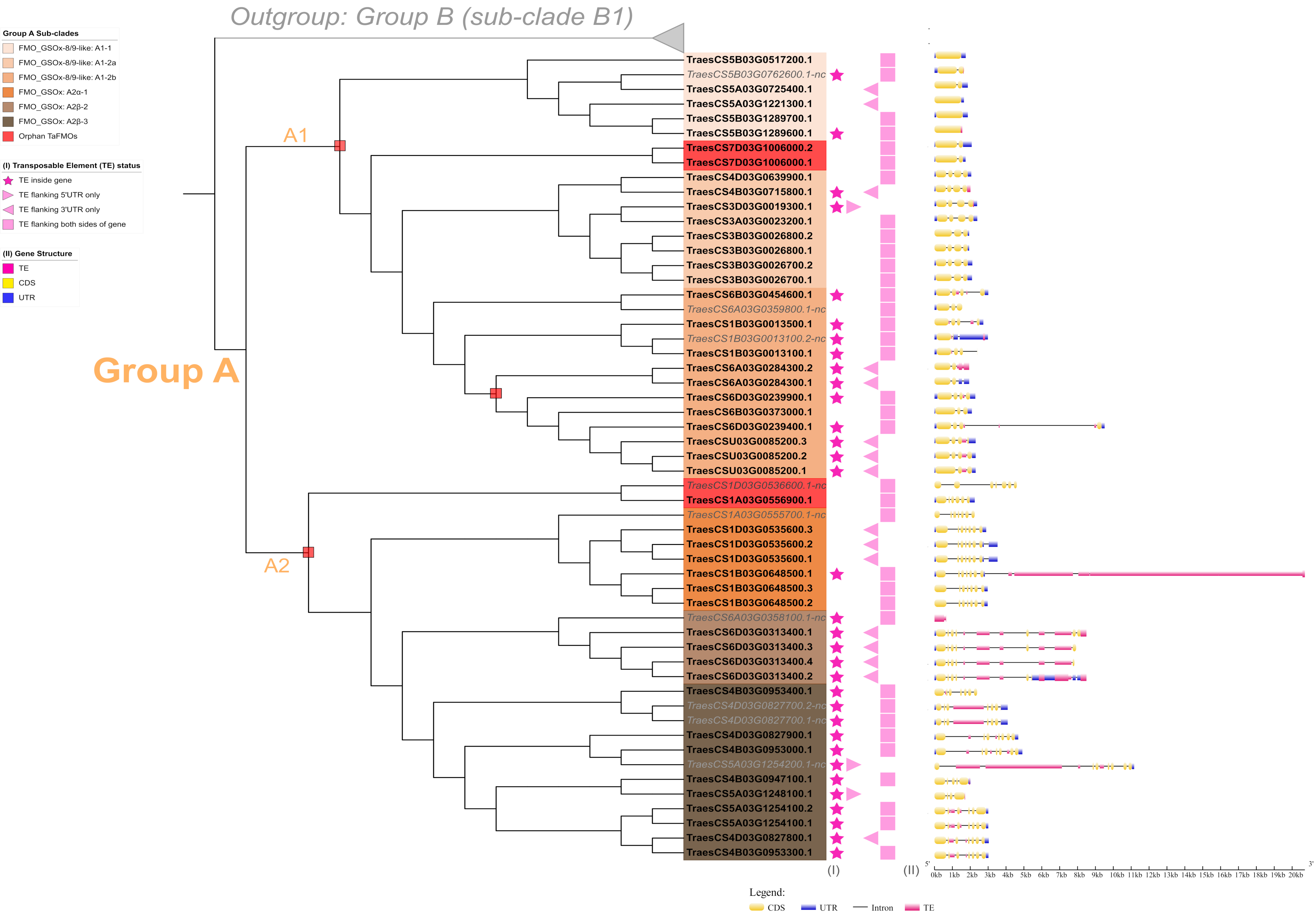

Outgroup: Group B (sub-clade B1)
Group A Sub-clades
FMO_GSOx-8/9-like: A1-1
FMO_GSOx-8/9-like: A1-2a
FMO_GSOx-8/9-like: A1-2b
FMO_GSOx: A2α-1
FMO_GSOx: A2β-2
FMO_GSOx: A2β-3
Orphan TaFMOs
(I) Transposable Element (TE) status
TE inside gene
TE flanking 5'UTR only
TE flanking 3'UTR only
TE flanking both sides of gene
(II) Gene Structure
TE
CDS
UTR
Group A
A1
A2
TraesCS5B03G0517200.1
TraesCS5B03G0762600.1-nc
TraesCS5A03G0725400.1
TraesCS5A03G1221300.1
TraesCS5B03G1289700.1
TraesCS5B03G1289600.1
TraesCS7D03G1006000.2
TraesCS7D03G1006000.1
TraesCS4D03G0639900.1
TraesCS4B03G0715800.1
TraesCS3D03G0019300.1
TraesCS3A03G0023200.1
TraesCS3B03G0026800.2
TraesCS3B03G0026800.1
TraesCS3B03G0026700.2
TraesCS3B03G0026700.1
TraesCS6B03G0454600.1
TraesCS6A03G0359800.1-nc
TraesCS1B03G0013500.1
TraesCS1B03G0013100.2-nc
TraesCS1B03G0013100.1
TraesCS6A03G0284300.2
TraesCS6A03G0284300.1
TraesCS6D03G0239900.1
TraesCS6B03G0373000.1
TraesCS6D03G0239400.1
TraesCSU03G0085200.3
TraesCSU03G0085200.2
TraesCSU03G0085200.1
TraesCS1D03G0536600.1-nc
TraesCS1A03G0556900.1
TraesCS1A03G0555700.1-nc
TraesCS1D03G0535600.3
TraesCS1D03G0535600.2
TraesCS1D03G0535600.1
TraesCS1B03G0648500.1
TraesCS1B03G0648500.3
TraesCS1B03G0648500.2
TraesCS6A03G0358100.1-nc
TraesCS6D03G0313400.1
TraesCS6D03G0313400.3
TraesCS6D03G0313400.4
TraesCS6D03G0313400.2
TraesCS4B03G0953400.1
TraesCS4D03G0827700.2-nc
TraesCS4D03G0827700.1-nc
TraesCS4D03G0827900.1
TraesCS4B03G0953000.1
TraesCS5A03G1254200.1-nc
TraesCS4B03G0947100.1
TraesCS5A03G1248100.1
TraesCS5A03G1254100.2
TraesCS5A03G1254100.1
TraesCS4D03G0827800.1
TraesCS4B03G0953300.1
(I)
(II)
5'
3'
0kb 1kb 2kb 3kb 4kb 5kb 6kb 7kb 8kb 9kb 10kb 11kb 12kb 13kb 14kb 15kb 16kb 17kb 18kb 19kb 20kb
Legend:
CDS
UTR
Intron
TE

**Figure S10** Graphical display of the gene structure and transposable element analysis for *TaFMOs* in Group A. Polytomy regions are denoted by a red box. The status of TE distribution is described in (I) for star: TE insertion inside gene; triangle: TE flanking one UTR of gene; and square: TE flanking both sides of gene. A 2 kb region upstream of the start codon and downstream of the stop codon were scanned for TEs. (II) The gene structure is displayed for each *TaFMO,* with graphical depiction of TE insertion, if present. The X-axis scale describes the size in kb of nucleotides for each region. For Group A, the number of exons present varies, from one or two (A1-1, orphan *TraesCS7D03G1006000*), two to four (A1-2a, A1-2b), and four to eight (A2). Group A TaFMOs have TE interruptions inside introns closer to the 3'UTR or inside the 3'UTR, with A1-2b, A2β-2 and A2β-3 containing the most TE insertions. The largest disruption seen in Group A occurs in *TraesCS1B03G0648500.1* inside the 3'UTR region, resulting in a total gene length of 20.71 kb.

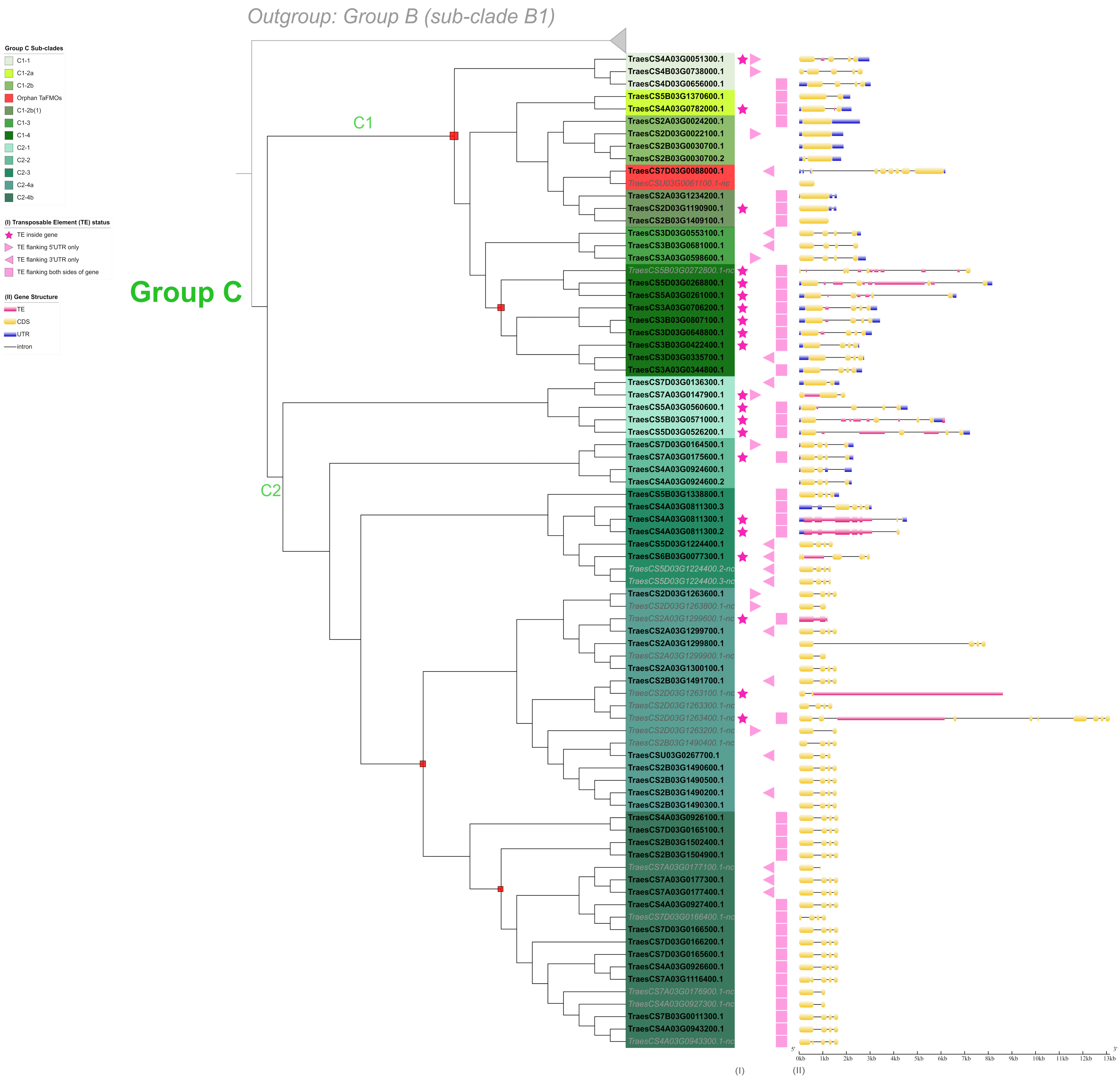
Outgroup: Group B (sub-clade B1)
Group C Sub-clades
C1-1
C1-2a
C1-2b
Orphan TaFMOs
C1-2b(1)
C1-3
C1-4
C2-1
C2-2
C2-3
C2-4a
C2-4b
(I) Transposable Element (TE) status
TE inside gene
TE flanking 5'UTR only
TE flanking 3'UTR only
TE flanking both sides of gene
(II) Gene Structure
TE
CDS
UTR
intron
Group C
C1
C2
TraesCS4A03G0051300.1
TraesCS4B03G0738000.1
TraesCS4D03G0656000.1
TraesCS5B03G1370600.1
TraesCS4A03G0782000.1
TraesCS2A03G0024200.1
TraesCS2D03G0022100.1
TraesCS2B03G0030700.1
TraesCS2B03G0030700.2
TraesCS7D03G0088000.1
TraesCSU03G0061100.1-nc
TraesCS2A03G1234200.1
TraesCS2D03G1190900.1
TraesCS2B03G1409100.1
TraesCS3D03G0553100.1
TraesCS3B03G0681000.1
TraesCS3A03G0598600.1
TraesCS5B03G0272800.1-nc
TraesCS5D03G0268800.1
TraesCS5A03G0261000.1
TraesCS3A03G0706200.1
TraesCS3B03G0807100.1
TraesCS3D03G0648800.1
TraesCS3B03G0422400.1
TraesCS3D03G0335700.1
TraesCS3A03G0344800.1
TraesCS7D03G0136300.1
TraesCS7A03G0147900.1
TraesCS5A03G0560600.1
TraesCS5B03G0571000.1
TraesCS5D03G0526200.1
TraesCS7D03G0164500.1
TraesCS7A03G0175600.1
TraesCS4A03G0924600.1
TraesCS4A03G0924600.2
TraesCS5B03G1338800.1
TraesCS4A03G0811300.3
TraesCS4A03G0811300.1
TraesCS4A03G0811300.2
TraesCS5D03G1224400.1
TraesCS6B03G0077300.1
TraesCS5D03G1224400.2-nc
TraesCS5D03G1224400.3-nc
TraesCS2D03G1263600.1
TraesCS2D03G1263800.1-nc
TraesCS2A03G1299600.1-nc
TraesCS2A03G1299700.1
TraesCS2A03G1299800.1
TraesCS2A03G1299900.1-nc
TraesCS2A03G1300100.1
TraesCS2B03G1491700.1
TraesCS2D03G1263100.1-nc
TraesCS2D03G1263300.1-nc
TraesCS2D03G1263400.1-nc
TraesCS2D03G1263200.1-nc
TraesCS2B03G1490400.1-nc
TraesCSU03G0267700.1
TraesCS2B03G1490600.1
TraesCS2B03G1490500.1
TraesCS2B03G1490200.1
TraesCS2B03G1490300.1
TraesCS4A03G0926100.1
TraesCS7D03G0165100.1
TraesCS2B03G1502400.1
TraesCS2B03G1504900.1
TraesCS7A03G0177100.1-nc
TraesCS7A03G0177300.1
TraesCS7A03G0177400.1
TraesCS4A03G0927400.1
TraesCS7D03G0166400.1-nc
TraesCS7D03G0166500.1
TraesCS7D03G0166200.1
TraesCS7D03G0165600.1
TraesCS4A03G0926600.1
TraesCS7A03G1116400.1
TraesCS7A03G0176900.1-nc
TraesCS4A03G0927300.1-nc
TraesCS7B03G0011300.1
TraesCS4A03G0943200.1
TraesCS4A03G0943300.1-nc
5'
3'
0kb
1kb
2kb
3kb
4kb
5kb
6kb
7kb
8kb
9kb
10kb
11kb
12kb
13kb
(I)
(II)

**Figure S11** Graphical display of the gene structure and transposable element analysis for *TaFMOs* in Group C. Polytomy regions are denoted by a red box. The status of TE distribution is described in (I) for star: TE insertion inside gene; triangle: TE flanking one UTR of gene; and square: TE flanking both sides of gene. A 2 kb region upstream of the start codon and downstream of the stop codon were scanned for TEs. (II) The gene structure is displayed for each *TaFMO*, with graphical depiction of TE insertion, if present. The X-axis scale describes the size in kb of nucleotides for each region. In Group C, exon numbers vary from one (C1-2b), or two to five (all rest of C), with seven exons present in the protein kinase receptor fusion FMO (*TraesCS7D03G0088000.1*).
